# Pharmacological or genetic inhibition of *Scn9a* protects human and mouse beta cells while dampening insulin secretion in type 1 diabetes

**DOI:** 10.1101/2023.06.11.544521

**Authors:** Peter Overby, Abby G. Gordon, Xiao-Qing Dai, Natalie S. Nahirney, Sophia Provenzano, WenQing Grace Sun, Yi Han Xia, Jiashuo Aaron Zhang, Evgeniy Panzhinskiy, Chieh Min Jamie Chu, Howard Haoning Cen, Søs Skovsø, Jelena Kolic, Andrew G. Edwards, Patrick E. MacDonald, James D. Johnson

## Abstract

Pancreatic β cells are essential for glucose homeostasis and are progressively lost during the development of type 1 diabetes. We previously demonstrated that the use-dependent Na^+^ channel inhibitor, carbamazepine, protects mouse β cells *in vitro* and *in vivo*. Here, we confirmed the protective effects of carbamazepine and other Na^+^ channel inhibitors in human β cells and investigated the specific role of the Na^+^ channel α subunit gene *Scn9a* (Nav1.7) in β cell function and survival. We generated β cell-specific knockout mice on the non-obese diabetic (NOD) background both *Ins1*Cre knock-in and AAV8-*Ins1*-Cre approaches resulting in significant reduction of β cell Na^+^ currents. Ca^2+^ responses and insulin secretion were significantly reduced, but only under the highest glucose conditions. Notably, carbamazepine treatment did not further alter insulin secretion or β cell survival in *Scn9a*-knockout islets, indicating that β cell *Scn9a* primarily mediates this drug’s measured effects. Consistent with this, β cell-specific deletion of *Scn9a* using AAV8-*Ins1*-Cre significantly reduced diabetes incidence in NOD mice. scRNAseq showed that this protection was associated with reduced *Ins2* and increased *Cdk8* in β cells. Collectively, our data show that *Scn9a* plays important roles in β cell excitability and survival during type 1 diabetes, thereby supporting this ion channel as a potential therapeutic target for β cell preservation.

**Article Highlights:**

- Hyperactivity has been proposed as a driver of β cell dysfunction and death in type 1 diabetes, but the specific role of Na^+^ currents has not been addressed.
- Targeted *Scn9a* deletion in β cells using *Ins1*^Cre^ knock-in mice reduced Na^+^ channel currents, reduced insulin release at high glucose concentrations, and protected β cells from apoptosis.
- Targeted deletion of *Scn9a* in β cells at 6 weeks of age using AAV8-*Ins1*Cre reduced diabetes incidence in NOD mice.
- These findings support the *Scn9a* Na^+^ channel as a novel therapeutic target in type 1 diabetes.

## Introduction

In type 1 diabetes, insulin secreting pancreatic β cells are mostly destroyed by autoimmune attack which leads to hyperglycemia and devastating disease complications (1). Protecting β cells from stress may therefore have therapeutic benefit (2), but there are few potential approaches that directly promote β cell survival. Excessive electrical activity, known as excitotoxicity, has been linked with increased cell death and dysfunction in multiple cell types, including β cells (3; 4). Insulin is a primary autoantigen both in murine and human type 1 diabetes pathogenesis (5; 6) and β cell hyper-activity may increase neo-autoantigen production, initiating and/or accelerating the disease (7). Excitotoxicity as a therapeutic target for early intervention in type 1 diabetes has been validated in pre-clinical and clinical research on verapamil, a voltage-gated Ca^2+^ channel inhibitor that prevents diabetes in mouse models (8) and delays diabetes progression in human RCTs (9–11). The effects of verapamil diminish over time, emphasizing the need to identify more drugs that can protect β cells from excitotoxicity to prevent or treat diabetes.

Voltage-gated Na^+^ channels have not been studied to the same extent as Ca^2+^ channels, and their physiological roles in β cells remain controversial despite Na^+^ currents being prominent in β cells from human and other mammals (12–14). In mouse β cells, Na^+^ channels have been reported to stay inactivated at resting membrane potentials and are therefore not believed to contribute to insulin secretion at physiological glucose concentrations (14–17). The Na^+^ channel blocker tetrodotoxin partially reduced glucose-induced insulin secretion in rat β cells, especially at high glucose concentrations (18). Previously, we used high-content screening to identify carbamazepine, a use-dependent Na^+^ channel inhibitor that is FDA-approved to treat epilepsy and other neurological conditions, as a drug that protects β cells (19). These results were validated in NOD mice where carbamazepine significantly reduced diabetes incidence (20).

*Scn9a* (Nav1.7) is the most prominent Na^+^ channel isoform expressed in both mouse and human β cells (21). Here, we examined the specific role of *Scn9a* in NOD mouse β cells and its contribution to type 1 diabetes initiation. Using complementary human islet and NOD mouse models, we confirmed that *Scn9a* is required for maximum insulin release in response to high glucose, and that *Scn9a* mediates the β cell protective effects of carbamazepine. The anti-diabetic effects of carbamazepine were phenocopied in inducible β cell-specific *Scn9a* knockout NOD mice. Together, these data suggest that Na^+^ channel-mediated excitotoxicity contributes significantly to type 1 diabetes pathophysiology.

## Research Design and Methods

### Human islets

Human islets from cadaveric donors were obtained from Alberta IsletCore(22), with research consent of donors and their families and approval by the Human Research Ethics Board at the University of Alberta (Pro00013094). Human islet use was approved by the ethical board from the UBC Clinical Research Ethics Board (H13-01865). The following donor numbers where used: R317, R318, R322 and R395 for islet cell survival assays, and R396, R398, R399, R406 and R407 for dynamic insulin secretion measured with islet perifusion *in vitro.* Full donor and isolation information can be found at www.isletcore.ca and the function and molecular features of these islets are on humanislets.com (23).

### Islet isolations and dispersion

Mice were euthanized by CO_2_, pancreata were inflated through ductal injection then incubated with collagenase. This was followed by filtration and hand picking of islets(24). Islets where either used whole or dispersed 24 hours post-isolation with 4 washes of Ca^2+^/Mg^2+^-free Minimal Essential Medium (Cellgro #15-015-CV) followed by gentle trypsinization (0.01%) by repeatedly pipetting up and down 30 times and resuspension in RPMI1640 (Thermo Fisher #11875-093 with 10% FBS, 1% PS)(25).

### Genetic manipulation in mice

All animal procedures were in accordance with the Canadian Council for Animal Care guidelines. All animal studies and protocols were approved by the UBC Animal Care Committee. Mice were housed in the Centre for Disease Modelling specific pathogen-free facility on a standard 12-hour light/12-hour dark cycle with ad libitum access to chow diet (PicoLab, Mouse Diet 20-5058). To generate NOD.*Ins1*^Cre/WT^, we contracted Jackson Laboratories to backcross B6(Cg)-Ins1tm1.1(cre)Thor/J(26) (Jackson Laboratory, No. 026801) > 12 times (27). NOD.*Scn9a*^flox/flox^ mice were generated on the NOD/ShiLtJ background by the Jackson Laboratory. We backcrossed/refreshed each parent colony every 5th generation. Female *Scn9a*^flox/wt^;*Ins1*^wt/wt^ and male *Scn9a*^flox/wt^;*Ins1*^cre/cre^ mice were set up as breeders, at age 8 to 9 weeks, to generate experimental mice. Females from the parental colonies were included only once as breeders to eliminate the risk of onset of hyperglycemia during pregnancy and blood glucose measurements were taken at the time of weaning for each breeder female. Cohorts included in generation of Kaplan-Meier curves were monitored twice per week and used only to determine hyperglycemia onset incidence and body mass changes. Diabetes was defined as two consecutive blood glucose measurements greater than or equal to 16 mmol/L, or a single measurement greater than or equal to 22 mmol/L. Mice that developed diabetes were immediately euthanized. Cohorts generated for other *in vivo* and *in vitro* analyses were terminated at 12 (AAV8-*Ins1*-Cre) or 18 weeks (*Ins1*^Cre^), prior to diabetes. For inducible knockout, AAV8-*Ins1-*Cre adeno-associated virus carrying *Ins1* promoter-driven Cre recombinase (AAV8-*Ins1*-Cre; produced by VectorBuilder Inc., Chicago, USA)(28) was injected into the intraperitoneal cavity of 6–7-week-old mice at a dose of 1×10^12^ VGP.

### Na^+^ channel subunit isoform expression and electrophysiology

The Chan-Zuckerberg Biohub’s CELLxGENE Discover Census with gget cellxgene module was used to query and analyze publicly available scRNA data (cellxgene.cziscience.com)(29). Genes of interest were selected from the voltage-gated channel Na^+^ α subunit gene group from the HGNC database (www.genenames.org). Human and mouse (*mus musculus*) scRNA data from pancreas were obtained from census version 2023-05-15. Dot plots of mean expression of each gene and fraction of expressed cells in each group were plotted with sc.pl.dotplot() function for both human and mouse results. Analysis was done in GoogleColab environment unless otherwise specified.

Islets shipped by overnight courier to Edmonton in RPMI 1640 medium and dispersed into single cells on arrival and cultured in RPMI-1640 (11875, Gibco) with 11.1 mM glucose, 10 % FBS (12483, Gibco), 100 units/mL penicillin, and 100 mg/mL streptomycin (15140, Gibco) for 1-3 days (30). Cells were patch-clamped in the whole-cell voltage-clamp or perforated current-clamp configurations in a heated bath at 32-35°C and patch pipettes with resistances of 4-5 MOhm after polishing. Whole-cell capacitance was recorded with the Sine+DC lock-in function of a HEKA EPC10 amplifier and PatchMaster software (HEKA Electronics, Germany) as previously described (30). Exocytosis and Na^+^ channel currents were measured 1-2 minutes after obtaining the whole-cell configuration. Exocytotic responses were measured in response to ten 500 ms depolarizations to 0 mV from a holding potential of -70 mV, and Na^+^ channel currents were activated by the membrane potentials ranging from -120 to - 20 mV. The bath solution contained (in mM): 118 NaCl, 20 tetraethylammonium-Cl, 5.6 KCl, 1.2 MgCl_2_, 2.6 CaCl_2_, 5 HEPES, and 5 glucose (pH 7.4 with NaOH). The patch pipettes were filled with (in mM): 125 Cs-glutamate, 10 CsCl, 10 NaCl, 1 MgCl_2_, 0.05 EGTA, 5 HEPES, 0.1 cAMP and 3 MgATP (pH 7.15 with CsOH). Action potential measurements were performed in current-clamp mode of the perforated patch-clamp configuration. Bath solution contained (in mM): 140 NaCl, 3.6 KCl, 1.5 CaCl_2_, 0.5 MgSO_4_, 10 HEPES, 0.5 NaH_2_PO_4_, 5 NaHCO_3_, and indicated concentrations of glucose (pH 7.3 with NaOH). The patch pipettes were filled with (in mM): 76 K_2_SO_4_, 10 KCl, 10 NaCl, 1 MgCl_2_ and 5 HEPES (pH 7.25 with KOH), and backfilled with 0.24 mg/ml amphotericin B (Sigma, cat# a9528). Mouse β- and α-cells were distinguished by characteristic differences in the voltage-dependent inactivation of Na^+^ currents (31; 32). Data were analyzed using FitMaster (HEKA Electronics) and Prism (GraphPad, San Diego, USA).

### Dynamic insulin secretion

To measure the dynamics of insulin secretion, 65 human islets or 100 mouse islets were added to columns and perifused (0.4 L/min) with 3 mM glucose KRB solution initially for 60 minutes and then exposed to different concentrations of glucose as indicated in the figures and described previously(33). Samples were stored at −20°C and insulin secretion was measured using a mouse insulin radioimmunoassay kit (Millipore Cat# HI-14K).

### Calcium imaging

Islets where isolated and dispersed according to established protocol and seeded on poly-D-lysine-coated glass coverslips in a 200 µL media droplet for 5 hours followed by overnight incubation in 6 well cell culture plate with 2 mL media. Fura-2-AM (5 μM) was added 30 min prior to the start of the experiment. Coverslips were then washed in HEPES buffer (137 mM NaCl, 5.4 mM KCl, 0.25 mM Na_2_HPO_4_, 0.44 mM KH_2_PO_4_, 2 mM CaCl_2_, 4.2 mM NaHCO_3_, 3 mM glucose, 10 mM HEPES, pH 7.4) and placed in a chamber before imaging at 37°C on a Zeiss Axiovert 200M microscope with a 10x Alpha-PlanFluar (Zeiss, Germany), using a CoolSNAP HQ2 CCD camera (Photometrics, Tucson, USA). Fura-2 fluorescence ratio (F340/F380 excitation; emission 510 nm) was recorded every 10 seconds.

### Islet cell survival assay

Disperse islets were seeded onto 384-well plates and stained with 50 ng/mL Hoechst 33342 (Invitrogen) and 0.5 μg/mL propidium iodide (PI), concentrations that do not affect islet cell viability(19). RPMI 1640 medium (Invitrogen) was supplemented to 5 mM glucose for human islets/cells and 10 mM glucose for mouse islets/cells. Media contained 100 U/mL penicillin, 100 μg/mL streptomycin (Invitrogen), 10% vol/vol FBS (Invitrogen). After 2 hours of staining, cells were incubated with a cytokine cocktail (25 ng/mL tumor necrosis factor-α (TNF-α), 10 ng/mL interleukin-1β (IL-1β), and 10 ng/mL interferon-γ (IFN-γ); R&D Systems) or thapsigargin (Millipore sigma). Immediately after the addition of test drugs (see table for overview), the cells were imaged at 37 °C and 5% CO_2_. Images were captured every second hour for 72 hours using a robotic microscope (ImageXpress^MICRO^ XLS, Molecular Devices) and analyzed using the MetaXpress multi-wave cell scoring package (Molecular Devices).

### Metabolic physiology

For all physiological tests, littermates at 18 weeks of age were subjected to a 5-hour fast prior. Fasting bodyweight and basal blood glucose concentration were recorded. For intraperitoneal glucose tolerance tests (IPGTT) and *in vivo* insulin secretion tests, 20% glucose solution in saline was administered by intraperitoneal injection to each mouse based on body weight (2 g/kg). OneTouch glucose strips and a OneTouch Ultra2 glucose meter were used to measure glucose in blood drawn from the tail vein. Blood collections for *in vivo* insulin secretion and insulin tolerance test (ITT) were collected from the saphenous vein. To minimize stress, animals were shaved to expose the saphenous vein prior to the fast.

### Histology, immunofluorescence and islet infiltration scoring

Pancreata were fixed in 4% paraformaldehyde for 24 hours at 4°C, then transferred to 70% ethanol for storage. Samples were paraffin embedded, sectioned, then stained with either hematoxylin/eosin (H&E), hematoxylin alone, terminal deoxynucleotidyl transferase dUTP nick end labelling (TUNEL), or insulin antibodies, then imaged at 20X magnification by WaxIT Histology Services Inc. (Vancouver, Canada). Pancreatic β cell area was quantified by normalizing insulin positive area to the respective section size as determined by DAPI staining and was reported as a ratio of the two numbers. Additionally, insulin stain was used to determine insulitis proportion independent of H&E stain. To determine insulitis proportion, positive area was identified, and each islet was annotated, including mononuclear cells. Insulitis proportion was determined for each islet annotation analysis by normalizing non-insulin area to insulin positive area. Insulin positive stain of the adjacent section was used to verify insulin positive area for determination of TUNEL-positive proportion, with the exception that mononuclear cells were excluded. Images were analyzed using Aperio ImageScope software v12.4.0.5043 (Leica Biosystems, Wetzlar, Germany).

For *Scn9a* and β cell proliferation immunofluorescence imaging, we used Raybiotech guinea pig anti-insulin antibody (cat#129-10332) was used at 1:500 dilution and Invitrogen *Scn9a* rabbit polyclonal antibody (PA5-77727) at 1:250 dilution. JacksonImmuno secondary antibodies donkey anti-rabbit FITC (1:200) (cat#711-095-152) and donkey anti-guinea pig Cy3 (1:600) (cat#706-165-148) were used to counterstain. Slides were imaged on a Zeiss Axiovert 200M microscope with 20x air objective with standardized exposure times for each channel. Intensity thresholds were standardized in SlideBook software (Intelligent Imaging Innovations, Denver, USA). Images were exported as 8-bit TIFFs and merged using ImageJ. To quantify proliferation imaging, we used anti-rabbit proliferating cell nuclear antigen (PCNA) (New England Biolabs, D3H8P) XP® antibody at 1:800 dilution. These images were taken on a Zeiss Axiovert 200M microscope with a 100x Alpha-PlanFluar NA 1.45 oil objective (Zeiss, Germany), using a CoolSNAP HQ2 CCD camera (Photometrics, Tucson, USA). Number of proliferating β cells was determined by PCNA positive nuclei completely surrounded by insulin staining.

### RNA sequencing and pathway analysis

Whole islets (100) were flash-frozen in liquid nitrogen and stored at −80 °C. RNA was isolated using RNeasy Mini Kit (Qiagen #74106) according to manufacturer’s instructions. Library preparation, RNA sequencing, and bioinformatics support were provided by the UBC Biomedical Research Centre Core Facility (Vancouver, BC, CA). Sample quality control was performed using the Agilent 2100 Bioanalyzer System (RNA Pico LabChip Kit). Sequencing was performed on the Illumina NextSeq 500 with paired end 42bp × 42bp reads, targeting a read depth of 60 million. Demultiplexed read sequences were then aligned to the reference sequence (UCSC mm10) using STAR aligner (v2.5.0b)(34). Differential expression analysis was analyzed using DESeq2 R package (35). Pathway enrichment was performed using the clusterProfiler R package with the Gene Ontology Molecular Function database (36).

Single-cell library preparation and barcoding were performed using the Evercode Whole Transcriptome v2 Kit (Parse Biosciences). Sequencing was performed on the Illumina NextSeq 2000, targeting a depth of approximately 40,000 reads per cell. FASTQ generation was performed using bcl2fastq (Illumina), followed by alignment to a custom UCSC mm10 genome modified to include the Cre recombinase transgene sequence (obtained from NCBI, RefSeq: NC_005856.1). Analysis used Parse pipeline v1.4.1 and Seurat v5.3.0. Cells with <2,000 genes, >13,000 genes, or <400,000 total RNA counts were filtered out. Following data integration and scaling, we performed principal component analysis (PCA) and uniform manifold approximation and projection (UMAP) dimensionality reduction. Cell clustering was performed using the FindNeighbors and FindClusters. Data visualization used DimPlot and plotVolcano. Differential expression analysis used FindMarkers and Wilcoxon rank-sum test (significance defined as adjusted *p* value <0.05 with log2 fold change >0.25). Pathway enrichment was performed using the clusterProfiler R package with the Gene Ontology Molecular Function (36).

### Statistics, data analysis, and data/resource availability

Data are expressed as mean ± standard error of the mean (SEM). Statistical analysis was performed using the Mann-Whitney test, one-way ANOVA, or repeated measures two-way ANOVA as indicated. Diabetes incidence was measured by Kaplan-Meier analysis and quantified by log-rank test. (GraphPad Prism, GraphPad Software Inc., La Jolla, USA). Unless otherwise stated, *p* values <0.05 were considered significant. Data and resources are available upon request to the corresponding author.

## Results

### Effects of Na^+^ channel modulators on cell death and insulin secretion in human islets

We previously discovered that carbamazepine improves survival of MIN6 cells and primary mouse islet β cells(20; 37). As a next step towards clinical translation, we tested whether the pro-survival effects of carbamazepine were translatable to human β cells and generalizable to other Na^+^ channel inhibitors. We used our high-throughput, multi-parameter, kinetic imaging platform (19) and dispersed human islet cells to compare the effects of various Na^+^ channel modulator drugs on survival in the presence a cytotoxic cytokine cocktail containing TNF-α, IL-1β, and IFN-γ, or a DMSO/vehicle control. We also tested the effects of each drug on ER stress-associated β cell death induced by the ER Ca^2+^ pump inhibitor thapsigargin(38). Carbamazepine protected human islet cells from both cytokine-induced and ER-stress-induced apoptosis in a dose-dependent manner (Fig. 1A). Oxcarbazepine, a structurally related analog of carbamazepine, demonstrated similar dose-dependent pro-survival effect on human islet cells in the presence of both thapsigargin and cytokines (Fig. 1B). Ambroxol and spider toxin ProTxII were only protective in the context of thapsigargin (Fig. 1C,D). Veratridine, an enhancer of voltage-dependent Na^+^ channel activity, exhibited the opposing effect by increasing cell death (Fig. 1E). Collectively, these data suggest a pro-survival class effect of Na^+^ channel modulation in human islet cells that mirrors our findings in rodent β cells (19).

**Figure 1.**
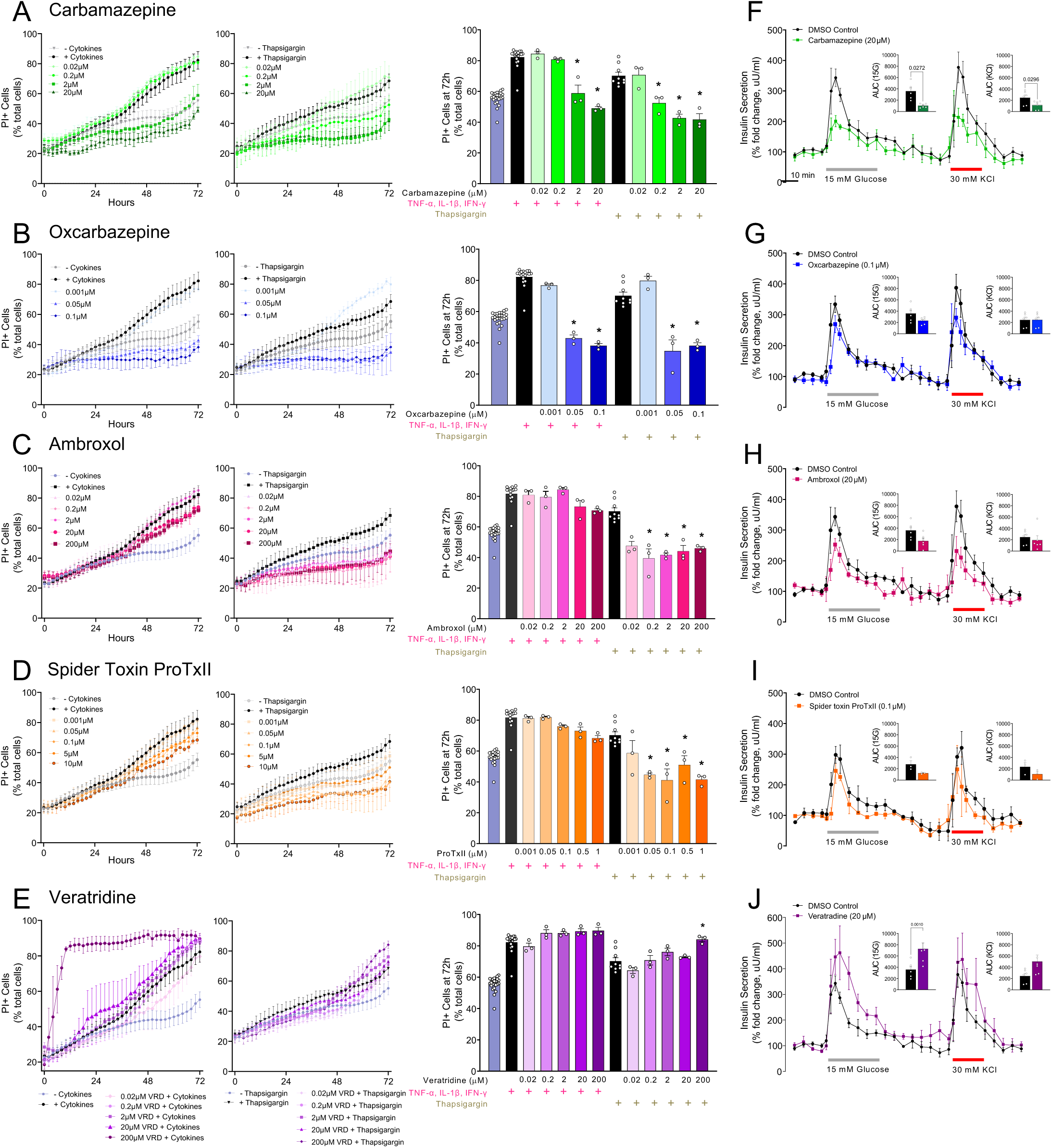
Na^+^ channel inhibitors reduced cytokine-induced cell death and have modest effects on insulin secretion in human islets. **(A-E)** Dispersed islet cells were seeded into 384-well plates and stained with Hoechst and propidium iodide (PI), then treated with either a cytokine cocktail or 1 μM thapsigargin, in combination with increasing concentrations of Na^+^ channel inhibitors: carbamazepine **(A)**, oxcarbazepine **(B)**, ambroxol **(C)**, spider toxin ProTxII **(D)**, or the Na^+^ channel enhancer veratridine **(E)**. Dimethyl sulfoxide (DMSO)-treated islets were used as the control condition, with the same results shown across all panels for consistency. Corresponding bar graphs show cell death at 72-hour time point (n = 3-18 human islet cell cultures, similar results were observed across human islet cultures from 4 donors). Cells were imaged with ImageXpress Micro and the percentage of PI-positive (PI+) cells was quantified. Statistical significance is indicated as *p<0.05. Data were evaluated using one-way ANOVA with Šídák’s multiple comparisons test, with significant defined relative to the cytokine-treated condition (Black). **(F-J)** 65 human islets were perifused with Krebs-Ringer bicarbonate buffer containing 15 mM glucose (grey line) then 30 mM KCl (red line) in combination with 20 μM carbamazepine **(F)**, 0.1 μM oxcarbazepine **(G)**, 20 μM ambroxol **(H)**, 0.1 μM spider toxin ProTxII **(I)**, or 20 μM veratridine **(J)**. Insets represent quantification of area under the curve (AUC) for the 15 mM glucose and 30 mM KCl phases (n = 3-6, mean ± SEM, p value shown). DMSO control represents the mean of corresponding controls from each specific run. AUC data were evaluated using 2-way ANOVA with multiple comparison (mixed models) and Dunnett’s multiple comparisons correction.

We also examined the effects of these Na^+^ channel-modulating drugs on glucose-stimulated insulin secretion from intact human islets. Carbamazepine was unique in its ability to significantly and selectively blunt the first phase of insulin secretion in response to 15 mM glucose or 30 mM KCl (Fig. 1F). Oxcarbazepine, ambroxol, and spider toxin ProTxII had modest, insignificant effects on insulin secretion under these conditions (Fig. 1G,H,I). Veratridine potentiated first and second phase insulin secretion at 15 mM glucose (Fig. 1J). These experiments show that, in general, protective Na^+^ channel modulating drugs tend to have modest inhibitory effects on insulin secretion.

### Expression of Na^+^ channel subunits in human and mouse islet cells

Multiple genes encode voltage-gated Na^+^ channels(39). Studies report that the Na^+^ channel isoform with the highest expression in β cells is encoded by the gene *Scn9a* (Nav1.7)(19; 40; 41), with *Scn9a* accounting for ∼80% of total Na^+^ channel mRNA in whole mouse islets (42). We further investigated the cell type-specific expression of Na^+^ α subunit genes using public C57Bl6 mouse and human single cell RNA sequencing datasets. In both species, *SNC9A*/*Scn9a* was found in all islet cell types and was the most frequently expressed of the Na^+^ channel genes in β cells (Fig. 2A,B). In NOD mouse β cells, *Scn9a* is also the most highly expressed subunit (e.g. GSE228219 (43); our new scRNAseq data). These data confirm that *Scn9a* is the major Na^+^ channel α subunit gene in β cells and illustrate the need to study cell-specific genetic manipulation of *Scn9a*.

**Figure 2.**
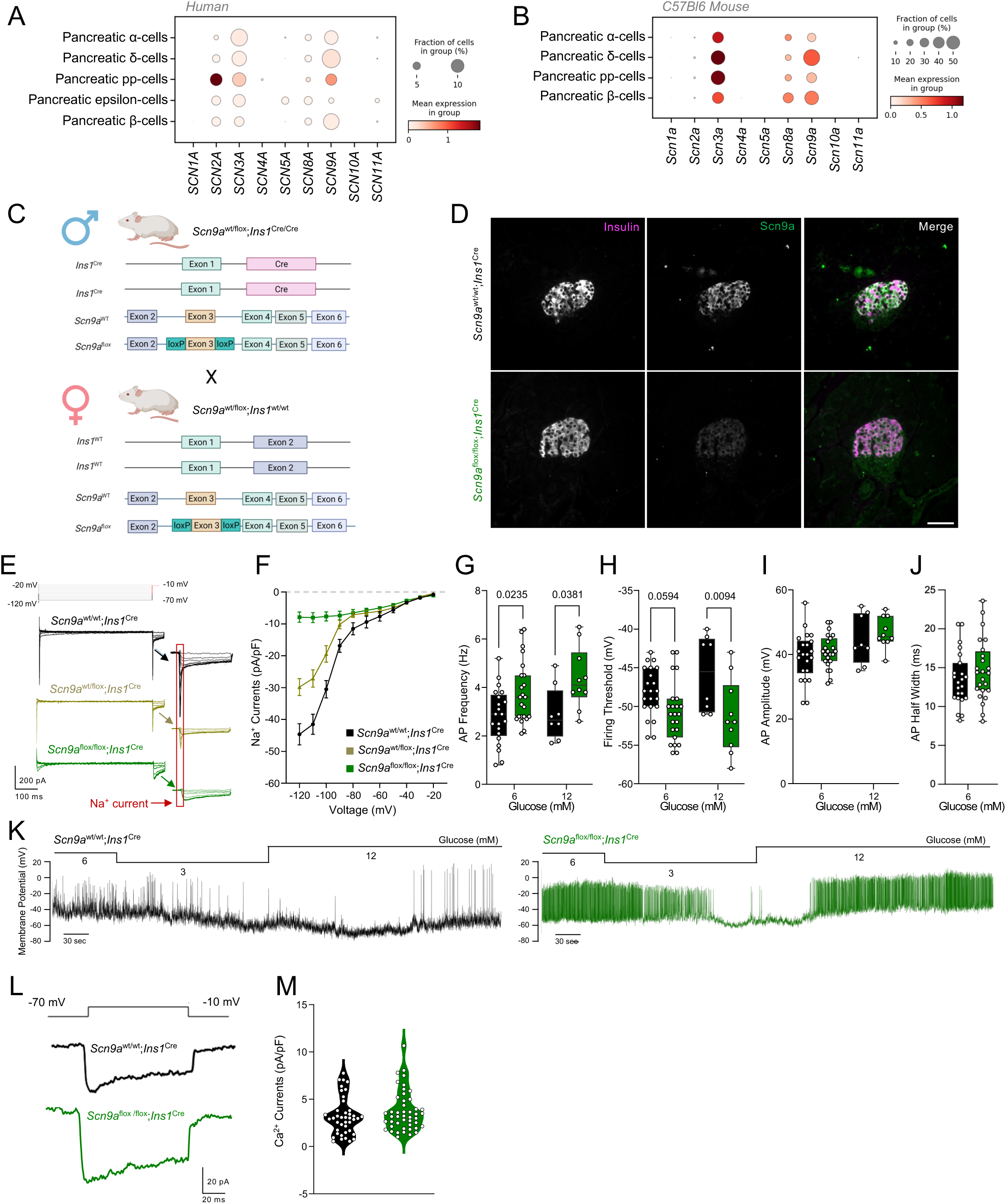
Voltage-gated Na^+^ channel alpha subunit expression and functional impact of *Scn9a* deletion in β cells. (**A,B)** Human and mouse islet cell scRNA data from CELLxGENE Discover Census. **(C)** Breeding strategy to produce β cell-specific *Scn9a* knockout mice and littermate controls. Created with Biorender.com. **(D)** Insulin (magenta) and Scn9a (green) immunostaining in NOD mouse pancreas. Scale bar is 20 mM. **(E)** Inward current traces, including Na^+^ current (red box), recorded from β cells with 2.6 mM Ca^2+^ in the extracellular solution. **(F)** Peak Na^+^ current normalized to cell size (pA/pF) as a function of voltage. **(G-J)** Action potential (AP) frequency, threshold, amplitude, and half width were measured during the 6 mM and 12 mM glucose periods in β cells. **(K)** Representative membrane potential recording from β cells exposed to 6, 3, and 12 mM glucose. Traces were obtained under the same experimental conditions used for analyses in (G-J). **(L,M)** Representative Ca^2+^ currents and quantification. Electrophysiology experiments were done in 18-week-old male mice; wildtype (black), heterozygous (brown), and knockout (green). Data were evaluated using unpaired t test.

### A cell-specific Scn9a knockout NOD mouse model with reduced β cell Na^+^ current

To study the specific roles of *Scn9a* in NOD mouse β cells, we generated *Scn9a*^flox/flox^;*Ins1*^Cre/wt^ (*Scn9a*^βKO^), *Scn9a*^flox/wt^;*Ins1*^Cre/wt^ (*Scn9a*^βHET^), and *Scn9a*^wt/wt^;*Ins1*^Cre/wt^ (*Scn9a*^WT^) littermates (Fig. 2C). Immunofluorescence imaging with a *Scn9a* antibody illustrated an expected reduction in fluorescence intensity in β cells from *Scn9a*^βKO^ mice (Fig. 2D). Critically, full *Scn9a*^βKO^ β cells exhibited a significant reduction in Na^+^ currents compared to *Scn9a*^WT^ β cells, while *Scn9a*^βHET^ β cells exhibited an intermediately reduced Na^+^ current across the voltage range (Fig. 2E,F). At potentials less than -80 mV, *Scn9a*^βKO^ and *Scn9a*^βHET^ β cells showed equivalent inhibition (Fig. 2F). These findings demonstrate a functional reduction in *Scn9a* activity in our model and support *Scn9a* as a major contributor to Na^+^ currents in NOD mouse β cells.

Next we assessed action potential behaviour in the perforated-patch, current-clamp configuration (Fig. 2G-K). The oscillatory patterns from *Scn9a*^βKO^ β cells were qualitatively distinguishable from *Scn9a*^WT^ β cells (Fig. 2K). Interestingly, quantified action potential frequency in *Scn9a*^βKO^ β cells was significantly increased relative to *Scn9a*^WT^ control β cells at both 6 mM and 12 mM glucose (Fig. 2G), while firing threshold was significantly reduced (Fig. 2H). However, there was no difference in action potential amplitude or half width (Fig. 2I,J). The paradoxical increased action potential firing may reflect compensatory changes occurring in our constitutive *Scn9a* knockout model. Consistent with this possibility, residual Na^+^ currents were observed in the -70 to -50 mV range in *Scn9a*^βKO^ β cells (Fig. 2E,F), which could reflect potential changes in the expression or activity of other voltage-gated Na+ channel. We were unable to identify up-regulation of *Scn3a* or other α subunit genes at the mRNA level (see below), but it remains possible that other Na^+^ channel proteins or co-factors could be upregulated in our knockout model. *Scn9a* deletion did not significantly change voltage-dependent Ca^2+^ currents in this model (Fig. 2L,M). Together, these data suggest that *Scn9a* Na^+^ channels contribute substantially to β cell Na^+^ currents and electrical activity in NOD mice, consistent with a previous report(41).

### Scn9a is required for high glucose response and mediates the effects of carbamazepine

We next assessed whether *Scn9a*-mediated Na^+^ currents contribute to insulin secretion in NOD mice. For this, we first used a sub-strain of NOD mice that is protected from diabetes due to the *Ins1* haploinsufficiency created by the Cre knock-in allele(27). Compared to wildtype littermate controls, high glucose-stimulated insulin secretion was reduced in islets isolated from both male and female *Scn9a*^βKO^ mice. This was significant at 15 mM glucose, but not at moderate glucose challenges (Fig. 3A,B). This agrees with published static incubation studies showing that insulin secretion response to 10 mM glucose was not different in islets from germline *Scn9a* knockouts, at least when compared with *Scn9a* heterozygotes(40). Importantly, the inhibitory effects of carbamazepine on 15 mM glucose-stimulated insulin secretion were absent in *Scn9a*^βKO^ mice (Fig. 3A,B), demonstrating that carbamazepine’s effect on insulin secretion is predominantly mediated through the *Scn9a* channel.

**Figure 3.**
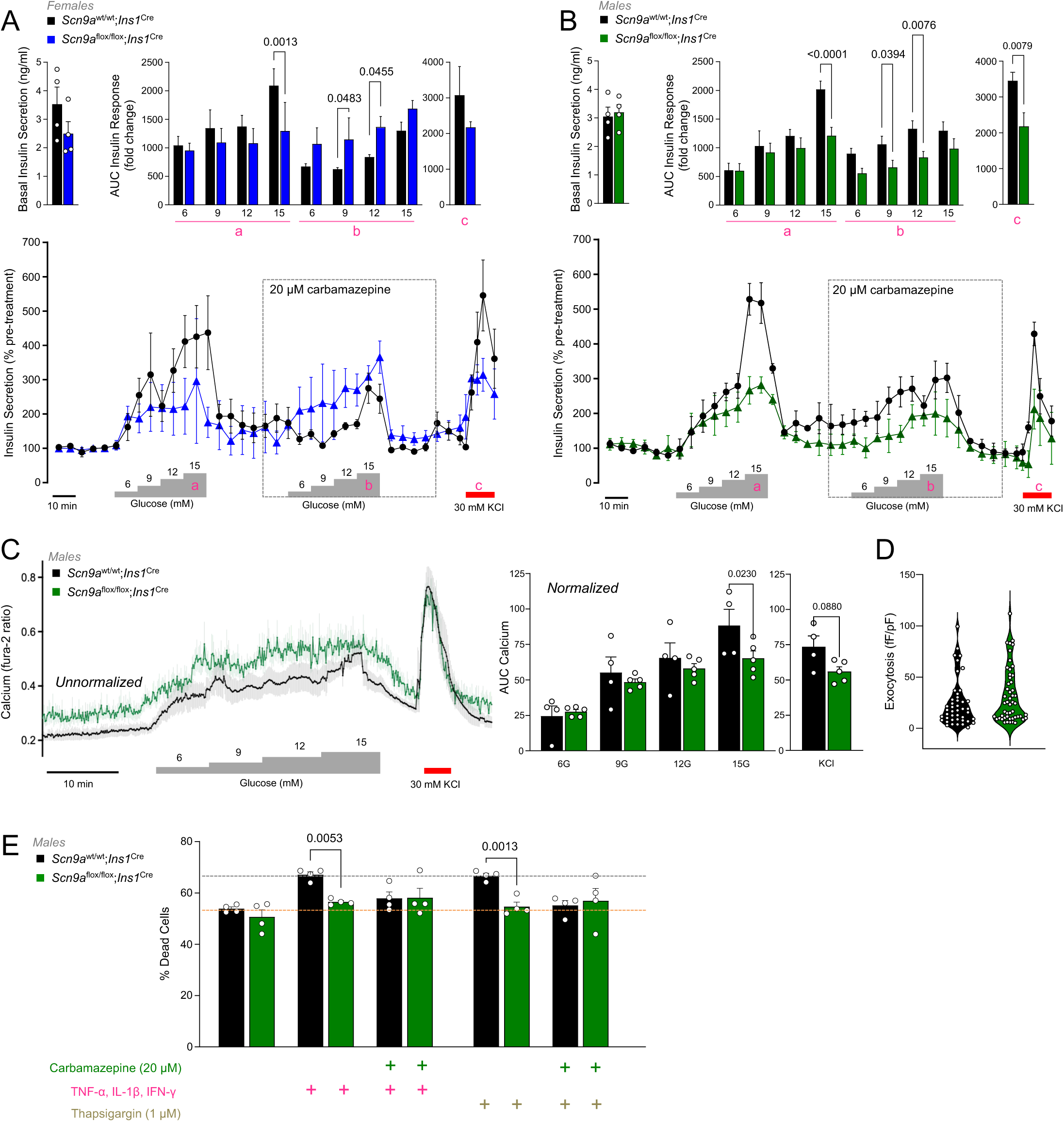
*Scn9a*-dependent regulation of glucose stimulated insulin secretion and protection by carbamazepine. **(A,B)** Glucose-dependent insulin secretion from 100 whole mouse islets, perifused with Krebs-Ringer bicarbonate buffer containing 6, 9, 12, and 15 mM glucose delivered in a ramp-like manner and repeated with or without 20 mM carbamazepine (indicated by dotted line -b) or 30 mM KCl (red line -c) (normalized to baseline at 3 mM glucose). Bar graphs above represent basal insulin secretion and quantification of area under the curve (AUC) of glucose ramp phase (a and b) or KCl (c). (n = 6). **(C)** Fura-2 ratio (340/380 nm) measured in dispersed islet cells perfused with Krebs-Ringer bicarbonate buffer containing 6, 9, 12, and 15 mM glucose delivered in a ramp-like manner (grey box) or 30 mM KCl (red line). Insets represent area under the curve (AUC) of KCl or individual glucose ramp phase. Each glucose ramp phase was normalized to baseline (3 mM glucose) (n = 4-5). AUC data were evaluated using 2-way ANOVA with multiple comparison (mixed models) with Dunnett correction for multiple comparisons. **(D)** Glucose-dependent exocytotic responses β cells measured as increases in cell membrane capacitance by whole-cell patch clamp at 5 mM glucose (n = 3-5). **(E)** Dispersed islet cells seeded onto 384-well plate were stained with Hoechst and PI. Cells were treated with a cytokine cocktail or 1 μM thapsigargin, in combination with either 20 μM carbamazepine or DMSO. The percentage of PI-positive (PI+) cells was calculated and represents quantification of cell death at 72 h time point (n = 4 islet cell culture from each of 4 mice). All experiments were done in 18-week-old mice; wildtype (black), female knockout (blue), and male knockout (green). Data were evaluated using 1-way ANOVA with Šídák’s multiple comparisons test.

To further define the molecular mechanisms of defective insulin secretion at 15 mM glucose in *Scn9a*^βKO^ cells, we loaded dispersed islet cells with Fura-2 to measure cytoplasmic Ca^2+^, which includes the contribution of multiple Ca^2+^ sources. In *Scn9a*^βKO^ cells, Ca^2+^ levels were elevated at baseline in 3 mM glucose (Fig. 3C). When normalized to baseline, we found a significant decrease in Ca^2+^ influx at 15 mM glucose exposure (Fig. 3C), consistent with the dynamic insulin secretion data above. Voltage-dependent increases in plasma membrane capacitance in patch-clamped β cells, which reflect the distal steps in exocytosis, were not different between *Scn9a*^βKO^ mice and controls (Fig. 3D). These findings indicate that the selective reduction in 15 mM glucose-stimulated insulin secretion is associated with reduced Ca^2+^ responsive from baseline, rather than a block of distal exocytotic machinery.

### Scn9a is required for stress-induced β cell death and mediates the effects of carbamazepine

We have established that pharmacological inhibition of Na^+^ channels protects MIN6 cells(19), primary mouse islets, and human islets (Fig. 1) from stresses relevant to type 1 diabetes. To test the hypothesis that β cell specific removal of the *Scn9a* channel protects primary murine β cells from cytokine-induced and thapsigargin-induced apoptosis, we employed the same multi-parameter imaging platform used to measure cell death in human islet cells above and dispersed cells from our *Scn9a*-deficient model. Cytokines and thapsigargin reproducibly induced death in primary islet cells from wildtype mice, whereas *Scn9a*^βKO^ islets were almost completely protected (Fig. 3E). We used cells from control and *Scn9a*^βKO^ mice to determine the contribution of this gene product to the pro-survival effects of carbamazepine(19). As expected, carbamazepine reduced cell death levels to those observed in DMSO controls (Fig. 3E). However, the addition of carbamazepine to *Scn9*a-deficient β cells did not yield a further reduction in cell death, suggesting that the pro-survival effects of carbamazepine are primarily mediated through the *Scn9a* channel. These data again point to the specificity of carbamazepine for β cell *Scn9a* and rule out major off-target effects.

### Diabetes incidence and glucose tolerance in β cells lacking Scn9a and Ins1 allele

Given the effects of *Scn9a* deletion on β cell survival *in vitro*, we next examined β cell turnover *in vivo*. While traditional categorical insulitis scoring was unable to identify significant differences between genotypes in male or female mice (Fig. 4A), continuous quantification of the ratio of insulin positive staining-to-insulitis in each islet identified reduced insulitis in male *Scn9a*^βKO^ mice when compared to littermate controls (Fig. 4B). We next stained pancreas sections from 18-week-old NOD mice for insulin and TUNEL or proliferating cell nuclear antigen (PCNA). Neither apoptosis nor proliferation were affected by β cell-specific *Scn9a* deletion (Fig. 4C,D). We also found no significant differences in insulin staining intensity (Fig. 4E) or islet size (Fig. 4F). We did however observe a statistically significant reduction in β cell area in male and female *Scn9a*^βKO^ mice relative to controls (Fig. 4G). We next examined islet insulin content because we have previously reported that carbamazepine increases insulin production (37) and isolated islets from global *Scn9a* knockout mice accumulate insulin protein in their β cells(37). However, there was no significant difference in acid/ethanol-extracted insulin protein content in *Scn9a*^βKO^ islets (Fig. 4H). These data point to a minor role for *Scn9a* in the maintenance of normal β cell mass and insulin production under non-diabetic conditions.

**Figure 4.**
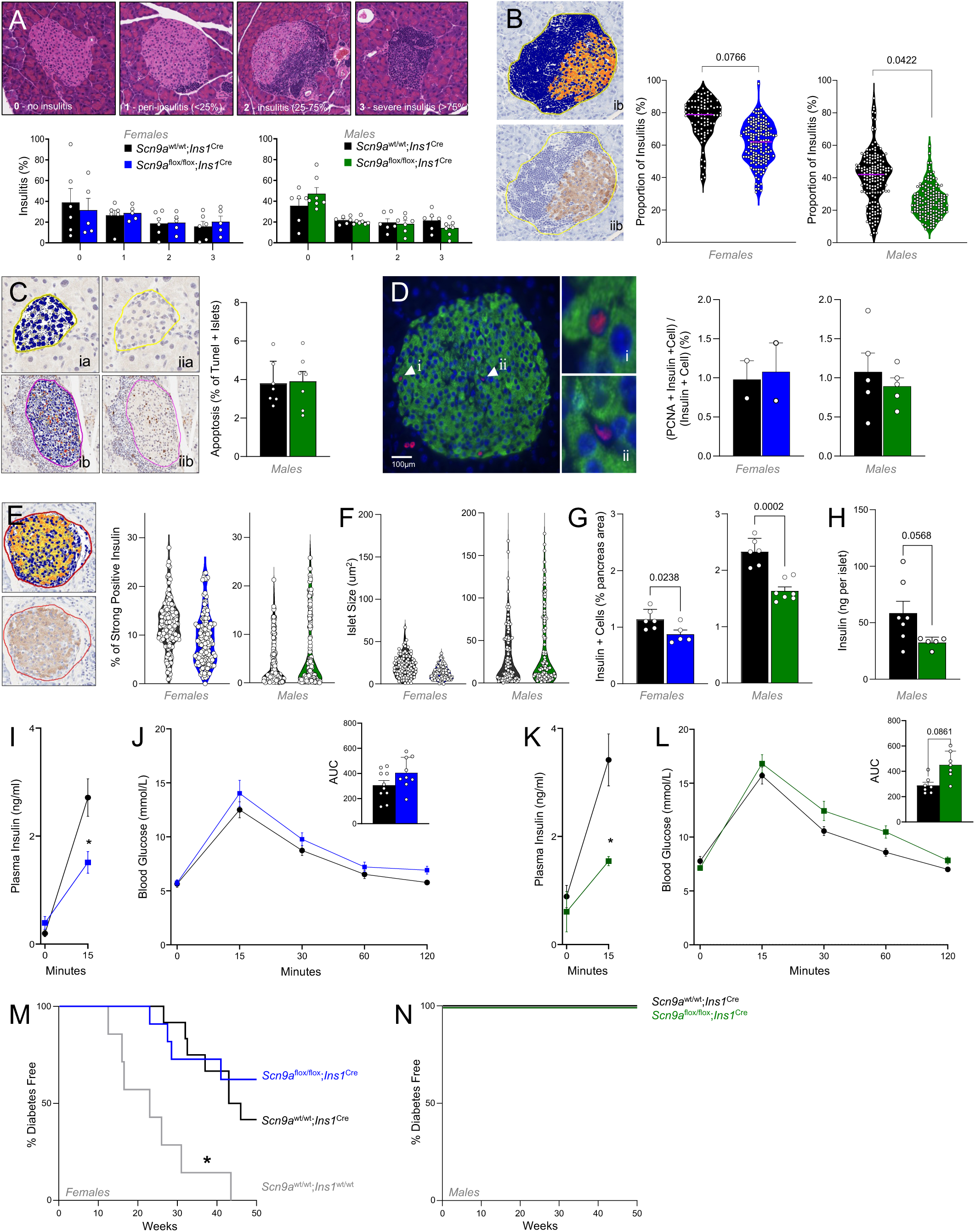
Life-long β cell *Scn9a* deletion reduced insulitis but impairs glucose stimulated insulin secretion and accelerates diabetes NOD mice. **(A)** Insulitis scoring using the common categorial method (0 –no insulitis, 1 –peri-insulitis with <25% peripheral immune invasion, 2 –insulitis with 25-75% immune invasion, 3 –severe insulitis with >75% immune invasion) in 18-week-old mice (n = 5-6 mice). **(B)** Insulitis scoring data using a continuous method (n = 5-6). Islets stained for insulin and islet periphery outlined in yellow. Violin plots show insulitis proportion (n = 6). **(C)** Quantification of apoptosis using TUNEL staining in pre-diabetic NOD mice (n = 7). **(D)** Quantification of β cell proliferation (i = proliferating β cell; ii = proliferating non-β cell) in pre-diabetic NOD mice (n = 4-5). **(E)** Insulin staining intensity and **(F)** islet size. **(G)** Quantification of β cell area (n = 5-7). **(H)** Insulin content in whole islets (n = 5-7). *p<0.05. Data were evaluated using Nested one-way ANOVA or One-way ANOVA. (I-L) Insulin secretion and glucose tolerance after intraperitoneal injection with 2 g/kg of glucose in 18-week-old NOD mice. Data were evaluated using one-way ANOVA. **(I,K)** Blood was collected at 0 (baseline) and 15 minutes for *in vivo* insulin secretion. **(J,L)** Blood glucose was measured as 0, 15, 30, 60 and 120 minutes after injection. Insets represent area under the curve (AUC) (n = 7-10). **(M,N)** Kaplan-Meier plot denoting diabetes incidence up to 50 weeks of age. (n = 12-13). Survival analysis was performed using Log-rank (Mantel-Cox) test *p<0.05.

Physiological analysis in 18-week-old male and female mice showed that *in vivo* insulin secretion 15 minutes after glucose challenge was significantly reduced in *Scn9a*^βKO^ mice relative to *Scn9a*^WT^ littermate controls (Fig. 4I,K). Glucose tolerance was comparable between groups (Fig. 4J,L), indicating that, while lower than controls, *Scn9a*^βKO^ mice have sufficient insulin capacity to maintain near-normal glucose homeostasis. These β cell-specific *Scn9a*^βKO^ mice were generated using the *Ins1*^Cre^ knock-in/replacement allele, which we have shown delays type 1 diabetes due to both the loss of the autoantigenic *Ins1* allele and presence of Cre itself (27). In this context, *Scn9a*^βKO^ did not offer further protection from type 1 diabetes, with incidence being statistically similar between female *Scn9a*^flox/flox^;*Ins1*^Cre^ and *Scn9a*^wt/wt^;*Ins1*^Cre^ NOD mice over 50 weeks (Fig. 4M). No male mice of either genotype developed diabetes (Fig. 4N). Therefore, we were precluded from testing whether *Scn9a* knockout protects this mouse model from type 1 diabetes.

### Transcriptomic analysis of whole Scn9a^βKO^ islets

To explore the molecular mechanisms mediating the effects of *Scn9a* deletion, we performed bulk RNA sequencing on whole islets isolated from male *Scn9a*^flox/flox^;*Ins1*^Cre^ and *Scn9a*^wt/wt^;*Ins1*^Cre^ NOD mice (Fig. 5A). We confirmed that *Scn9a* mRNA (exon 3) was significantly reduced in islets from *Scn9a*^βKO^ mice (3.5-fold decrease in islets that also contain non-β cells; Fig. 5B). Differential gene analysis identified 5 other significantly decreased mRNAs in *Scn9a*^βKO^ islets (Fig. 5C,D), including neuropeptide Y (*Npy*) which has been implicated in β cell function and survival (44; 45) and cytochrome P450-26B1 (*Cyp26b1*) which is involved in embryonic development and cell differentiation(46). Interestingly, Cyp26b1 degrades carbamazepine (47) and carbamazepine has a negative effect on cytochrome P450 enzyme activity and expression (48). The farnesoid X receptor (*Nr1h4*) has multiple roles in β cells (49). Seven in absentia homolog 2 (*Siah2*) is responsible for degradation of proteins and involved in cellular processes including cell cycle progression, DNA repair, apoptosis, and hypoxia response (50).

**Figure 5.**
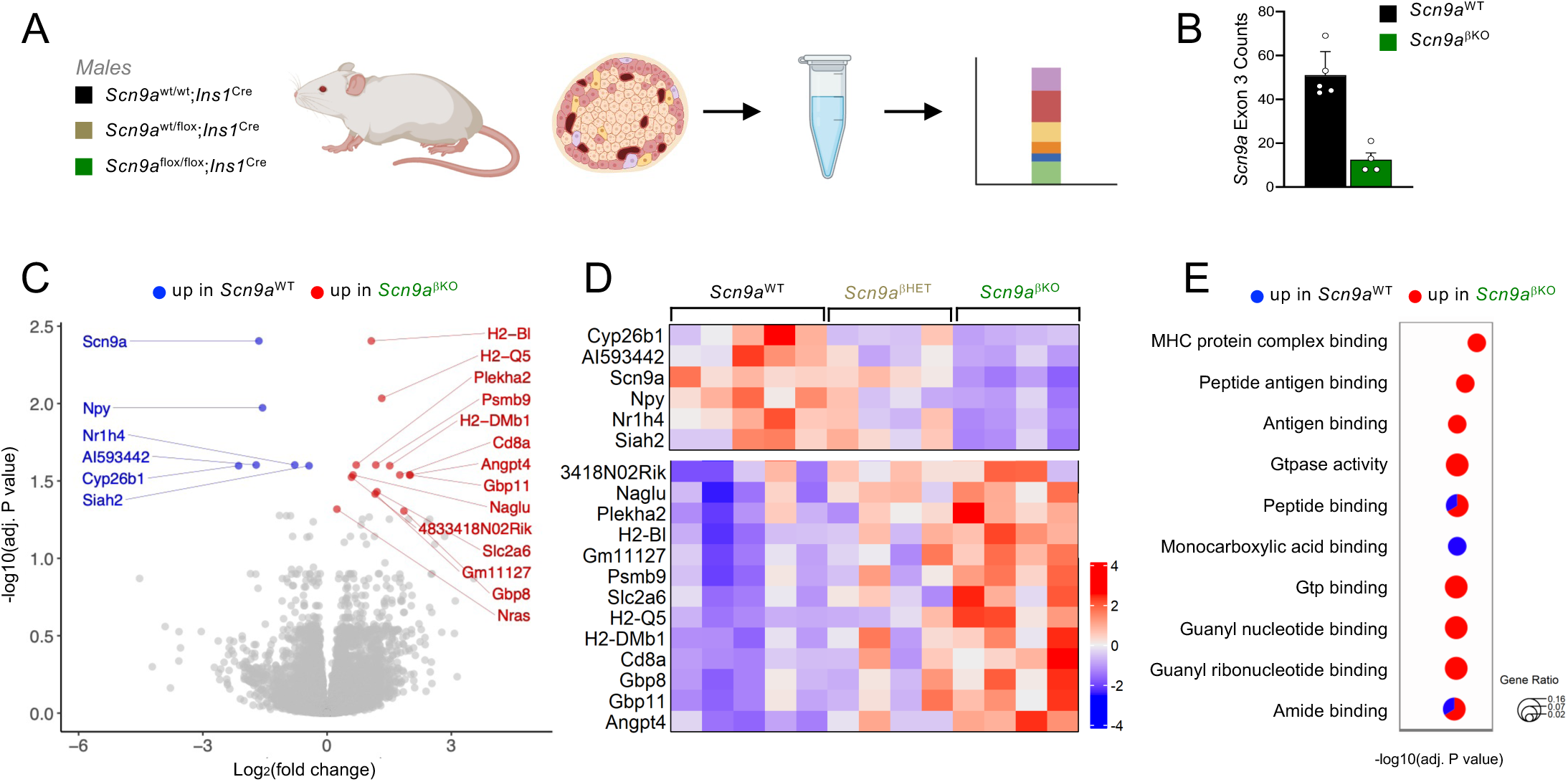
Bulk RNA sequencing data from male *Scn9a* KO NOD mouse islets show differentially expressed genes. **(A)** Study design shown as schematic diagram of experiments conducted on islets from wildtype (*Scn9a*^WT^), heterozygous (*Scn9a*^βHET^), and complete *Scn9a* knockout (*Scn9a*^βKO^) 18-week-old male NOD mice (n = 4-5). Created with Biorender.com. **(B)** Quantification of *Scn9a* exon 3 (floxed portion of the *Scn9a* gene) counts from RNA-seq analysis between genotypes. **(C)** Volcano plot showing differentially expressed mRNAs between WT and KO groups. **(D)** Heatmap showing average expression of differentially expressed genes, with values representing per sample expression grouped by genotype. **(E).** Top significantly enriched Reactome pathways from the top significantly differentially expressed genes, based on the Gene Ontology (GO) Molecular Function (MF) database.

We also identified 13 mRNAs that were significantly increased in *Scn9a*^βKO^ islets. Strikingly, multiple genes associated the major histocompatibility complex (MHC) class I and II were upregulated in islet cells from *Scn9a*^βKO^ mice (Fig. 5C,D), including H2-BI, H2-Q2, and H2-DMB1 (also known as H2-M; class II), which are involved in antigen presentation to CD8-positive T cells (51; 52). There was an increase in the CD8 alpha subunit mRNA in these islets. Both the H2-Q2 and H2-DMB1 genes have been implicated in the development of autoimmunity in NOD mice (53; 54) In accordance, we found the 3 top pathways enriched in this analysis, MHC protein complex binding, peptide antigen binding, and antigen binding, are all involved in the process of presenting antigens to the immune system (55)(Fig. 5E). Solute carrier family 2 member 6 (*Slc2a6*; aka GLUT6) was also upregulated in *Scn9a*^βKO^ islets. Interestingly, pancreatic islets isolated from GLUT6-deficient *Lep*^ob/ob^ mice secreted a greater amount of insulin in response to high levels of glucose compared to wildtype controls (56). Other upregulated mRNAs included: pleckstrin homology domain containing A2 (*Plekha2*), which binds specifically to phosphatidylinositol 3,4-diphosphate and localize proteins to the membrane (57); proteasome 20S subunit beta 9 (*Psmb9*), a component of the proteosome (58); N-acetyl-alpha-glucosaminidase (*Naglu*), an enzyme involved in the degradation of heparan sulfate (59); guanylate binding protein 6 (Gbp6/8/11), which is involved in cytokine signalling and confers protection from pathogens (60); NRAS proto-oncogene GTPase (*Nras*); angiopoietin 4 (*Angpt4*), which regulates the receptor tyrosine kinase TEK/TIE2 (61). These observations, on balance, suggest that the *Scn9a* channel in β cells could directly or indirectly modulate the immune system and provide further insight into the molecular mechanism of β cell protection in type 1 diabetes. Notably, we did not observe significant compensatory upregulation in other Na^+^ channel subunit genes at the mRNA level.

### AAV8Ins1Cre-mediated deletion of Scn9a reduces diabetes incidence in NOD mice

As observed in Fig. 4, NOD.*Scn9a*^flox^;*Ins1*^Cre^ mice were not suitable for studying the impact of *Scn9a* in type 1 diabetes progression (27). To knockout *Scn9a* specifically in adolescent β cells, while leaving the autoantigenic *Ins1* gene intact, we used the pancreas-biased AAV8 virus serotype and a highly specific *Ins1* promoter to deliver Cre for inducible deletion of *Scn9a* at 6-7 weeks (*Scn9a*^βiKO^)(Fig. 6A). This approach has comparable efficiency to Cre driver line models (28). First, we demonstrated high recombination efficiency and specificity with AAV8*Ins1*GFP in islets from wildtype NOD mice (Fig. 6B). Na^+^ current recordings confirmed that *Scn9a*^βiKO^ β cells had nearly complete loss of Na^+^ currents compared to *Scn9a*^WT^ β cells (Fig. 6C,D), with even less residual current than the life-long transgenic *Scn9a*^βKO^ model suggestive of less compensation. *Scn9a*^βiHET^ β cells showed an intermediate reduction across the range of holding potentials, although homozygous and heterozygous AAV8*Ins1*-mediated deletion showed similar current reduction at -80 mV (Fig. 6D).

**Figure 6.**
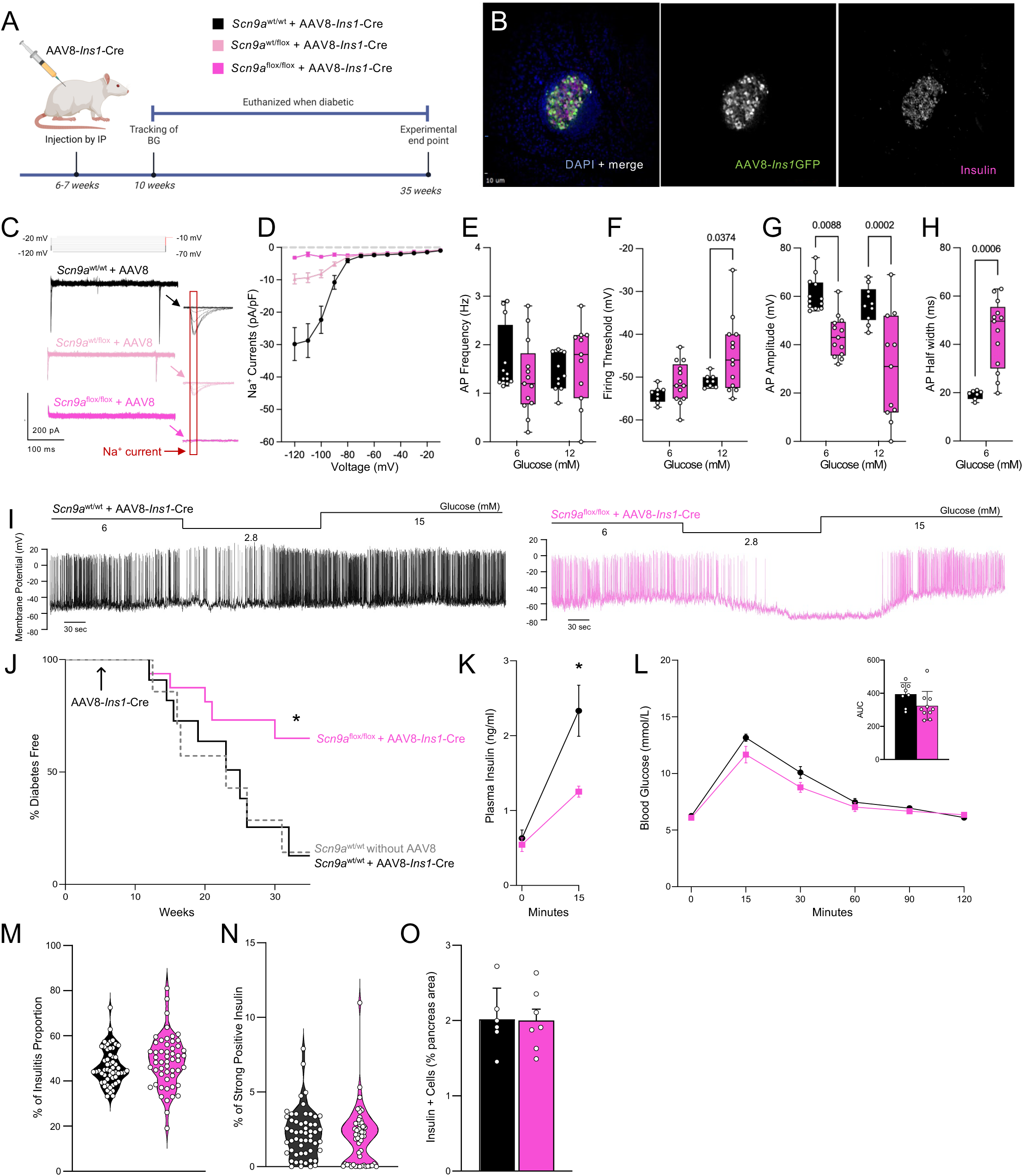
*Scn9a* deletion using AAV8-*Ins1*-Cre virus reduces Na^+^ currents in β cells and protects NOD mice from type 1 diabetes. **(A)** Schematic diagram of general study design for wildtype (black), heterozygous (light pink), and knockout (magenta) female NOD mice. Created with Biorender.com. **(B)** Efficiency of AAV8-*Ins1-*Cre virus in female NOD mice is illustrated using 1×10^12^ VGP of virus carrying GFP instead of Cre, imaged 4 weeks post injection. **(C)** Inward current traces, including Na^+^ current (red box), recorded from β cells with 2.6 mM Ca^2+^ in the extracellular solution. **(D)** Peak Na^+^ current normalized to cell size (pA/pF) as a function of voltage. **(E-H)** Action potential (AP) frequency, threshold, amplitude, and half width were measured during the 6 mM and 15 mM glucose periods in β cells. **(I)** Representative membrane potential recording from β cells exposed to 6, 2.8, and 15 mM glucose. Traces were obtained under the same experimental conditions used for analyses in (E-H). All electrophysiology experiments were done in 10-week-old female mice. Data were evaluated using unpaired t test. **(J)** Kaplan-Meier plot denoting diabetes incidence. Curves represent wildtype without AAV8 (grey dotted line, from *Scn9a*^wt/wt^;*Ins1*^wt/wt^ cohort shown in Fig. 4M), wildtype with AAV8 (black), and *Scn9a* knockout mice (pink) (n = 7-16). Mice received 1×10^12^ VGP AAV8-*Ins1*-Cre by intraperitoneal injection at 6-7 weeks, as indicated by arrow. Survival analysis was performed using Log-rank (Mantel-Cox) test *p<0.05. **(K,L)** *In vivo* insulin secretion and glucose tolerance tests following intraperitoneal injection with 2 g/kg of glucose in 12-week-old female mice. Data were evaluated using one-way ANOVA. **(K)** Blood was collected at 0 (baseline) and 15 minutes for *in vivo* insulin secretion. **(L)** Blood glucose was measured as 0, 15, 30, 60 and 120 minutes after injection. Inset represents area under the curve (AUC) (n = 9-10). **(M)** Violin plot showing the insulitis proportion in 12-week-old mice. **(N)** Insulin staining intensity quantified as a percentage of insulin positive area in 12-week-old mice (n = 6-7). **(O)** Insulin-positive cell area, as a percentage of total pancreas area, in 12-week-old mice (n = 6-7). *p<0.05.

Current clamp analysis of action potential behaviour, *Scn9a*^βiHET^ β cells differed from controls in a way that was distinct from the *Scn9a*^βiKO^ model (Fig. 6E-I). Unlike the *Scn9a* knockout mice generated with the *Ins1*^Cre^ knock-in, there were no significant differences in action potential frequency (Fig. 6E); in fact, *Scn9a*^βiKO^ β cells exhibited significantly higher firing thresholds in at 15 mM glucose (Fig. 6F). *Scn9a*^βiKO^ β cells action potentials had significantly decreased amplitude (Fig. 6G) and increased width (Fig. 6H). The near-complete loss of Na^+^ currents following inducible deletion, together with more expected modifications of action potential dynamics, validated the use of the AAV8*Ins1*Cre approach to examine the consequences of *Scn9a* deletion in β cells while minimizing potential developmental or long-term compensatory changes.

We hypothesized that AAV8-mediated β cell-specific *Scn9a* deletion in NOD mice would phenocopy the delay in diabetes seen in with carbamazepine treatment (20). Indeed, we observed a significant reduction in diabetes incidence between *Scn9a*^βiKO^ and control female mice (Fig. 6J). Again, in the pre-diabetic state we observed a significant reduction in glucose-stimulated insulin secretion *in vivo* (Fig. 6K), despite no significant difference in glucose tolerance (Fig. 6L). We did not observe differences in insulitis proportion, insulin positive area, or strong insulin positive cell proportion between *Scn9a*^βiKO^ and their controls (Fig. 6M-O). Collectively, our data demonstrated that β cell *Scn9a* deletion at 6-7 weeks using AAV8*Ins1*Cre can prevent 50% of diabetes cases in our NOD mouse population by 35 weeks of age.

### Single-cell transcriptomic analysis of virus-induced β cell specific Scn9a knockout mice

To assess cell type-specific effects that mediate this protection from type 1 diabetes, we performed single-cell RNA sequencing on dispersed islets isolated from female *Scn9a*^flox/flox^ and *Scn9a*^wt/wt^ NOD mice following AAV8*Ins1*Cre mediated deletion of *Scn9a* (Fig. 7A). Clustering analysis revealed 6 distinct cell populations which were labeled based on the expression of canonical marker genes (Fig. 7B). Alignment to a custom reference *Mus musculus* (mm10) genome containing the Cre recombinase transgene showed a strong enrichment of Cre expression in the β cell population (Fig. 7C,D).

**Figure 7.**
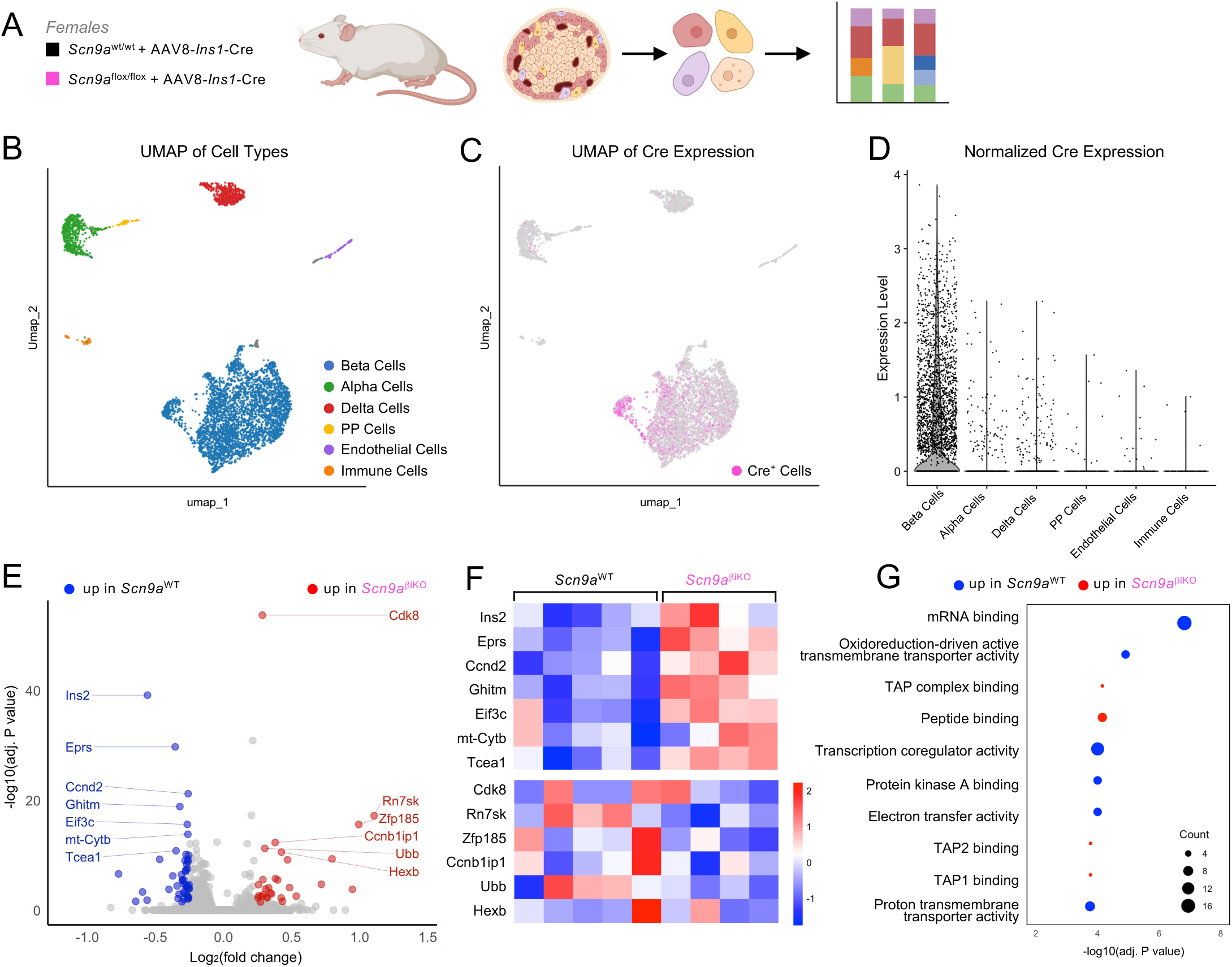
Single-cell RNA sequencing data from female *Scn9a* KO NOD mouse islets show differentially expressed genes. **(A)** Study design shown as schematic diagram of experiments conducted on islets from wildtype (*Scn9a*^WT^) and complete *Scn9a* knockout (*Scn9a*^βiKO^) 14-week-old female NOD mice (n = 4-5). Created with Biorender.com. **(B)** Uniform Manifold Approximation and projection (UMAP) of all single-cell transcriptomes, with clusters representing distinct cell populations based on canonical markers. **(C)** UMAP showing Cre-positive (Cre>0) cells across all identified clusters. **(D)** Normalized Cre expression levels across all identified clusters. **(E)** Volcano plot showing differentially expressed mRNAs between *Scn9a* WT and KO β cells. **(F)** Heatmap showing average expression of differentially expressed genes in β cells, with values representing per sample expression grouped by genotype. **(G).** Top significantly enriched Reactome pathways from the top significantly differentially expressed genes in β cells, based on the Gene Ontology (GO) Molecular Function (MF) database.

We identified 7 mRNAs that were significantly downregulated in *Scn9a*^βiKO^ β cells (Fig. 7E,F). The most significantly reduced gene was insulin 2 (*Ins2*), which carries the same autoantigenic B chain peptides as *Ins1* (6). Cyclin D2 (*Ccnd2*), an essential contributor to pancreatic β cell proliferation in mice (62), was also decreased. Other downregulated mRNAs relate to protein synthesis processes, including aminoacyl-tRNA synthetase (*Eprs*), eukaryotic initiation factor 3 subunit c (*Eif3c*), and transcription elongation factor A protein 1 (*Tcea1*). Several mitochondrial function genes were also downregulated, including growth hormone-inducible transmembrane protein (*Ghitm*), which helps maintain mitochondrial morphology (63), and mitochondrially encoded cytochrome B (*mt-Cytb*), a component of the electron transport chain (64).

We identified 6 mRNAs that were significantly upregulated in *Scn9a*^βiKO^ β cells (Fig. 7E,F), including cyclin-dependent kinase 8 (*Cdk8*), a key regulator of transcription with implications in insulin secretion and in modulating β cell apoptosis under oxidative stress (65), and RNA component of 7SK nuclear ribonucleoprotein (*Rn7sk*), which plays a critical role in regulating transcription elongation (66). Together, these data suggest that *Scn9a* loss broadly impacts transcriptional regulation in β cells.

Gene Ontology (GO) Molecular Function analysis of downregulated mRNAs revealed reduced activity in pathways associated with transcription and translation, including mRNA binding and transcription coregulator activity (Fig. 7G). In addition, mitochondrial-associated functions were decreased, including oxidoreduction-driven active transmembrane transporter activity, electron transfer activity, and proton transmembrane transporter activity. Consistent with bulk RNA sequencing analysis in Figure 5E, several top enriched pathways were associated with antigen processing, specifically with binding of the transporter associated with antigen processing (TAP) complex. This complex, composed of TAP1 and TAP2 subunits, facilitates the transport of cytosolic peptides into the endoplasmic reticulum for loading onto MHC Class I molecules (67). Variants in *TAP1* and *TAP2* have been associated with T1D susceptibility, with some variants linked to increased risk (68) and others reported to confer protection (69). In *Scn9a*^βiKO^ β cells, we did not observe significant compensatory upregulation in other Na^+^ channel subunit genes, although we did not have high statistical power to detect differences in weakly expressed mRNAs.

### Effects of heterozygous deletion of β cell Scn9a

In parallel, we studied β cells, islets, and mice that were heterozygous for the mutant *Scn9a* allele. Isolated β cells from *Scn9a* heterozygous mice had intermediately reduced Na^+^ currents across the range of holding potentials, but similar reduction at physiological voltages (Fig. 2E,F; Fig. 6C,D). Voltage-clamp showed that the action potential firing patterns of heterozygous *Scn9a-*deficient β cells were distinguishable from those seen in *Scn9a*^WT^ control mice (Fig. S1A), at 6 mM and 12 mM glucose, firing action potential, firing thresholds and potential height were unchanged (Fig. S1B). Cytosolic Ca^2+^ levels were elevated at baseline in 3 mM glucose in β cells from *Scn9a*^βHET^ mice (Fig. S1C). When normalized to baseline, we found a significant decrease in Ca^2+^ influx at 15 mM glucose exposure in *Scn9a*^βHET^ cells (Fig. S1C, *inset*), similar to *Scn9a*^βKO^ cells (Fig. 3C). Despite this, insulin secretion from *Scn9a*^βHET^ islets was more strongly inhibited than from *Scn9a*^βKO^ islets, with a near complete block of insulin secretion at all glucose concentrations (Fig. S1D,E) and inhibition of distal exocytosis mechanisms (Fig. S1F). In contrast to *Scn9a*^βKO^ mice, *Scn9a*^βHET^ mice demonstrated significantly worse glucose tolerance than their *Scn9a*^WT^ littermates (Fig. S2A-D). Type 1 diabetes had a rapid onset in life-long *Scn9a*^βHET^ females but was unchanged in males (Fig. S2E,F). AAV8-*Ins1-*Cre mediated removal of one *Scn9a* allele at 6-7 weeks also accelerated diabetes in female NOD mice (Fig. S3A,B,C). *Scn9a*^βHET^ mice from the *Ins1-*Cre cohort also exhibited increased insulitis (Fig. S3D). Mechanistically, *Scn9a*^βHET^ primary islet cells showed little to no protection from cytokine- or thapsigargin-induced death *in vitro* (Fig. S4A). In fact, we observed a significant increase in TUNEL positive cells, as well as a concomitant reduction in insulin area and insulin-positive cell number (Fig. S4B,C). Surprisingly, islet insulin content was increased in islets isolated from *Scn9a*^βHET^ mice relative to controls (Fig. S4D). Together, these data point to a complex effect of partial *Scn9a* loss in β cells but also provide clues regarding which aspects of the *Scn9a*^βKO^ phenotype are anti-diabetic.

## Discussion

The goal of the present study was to investigate the roles of Na^+^ channels and drugs that can modulate them in human and NOD mouse β cells. We found that multiple Na^+^ channel-modulators can protect human islet cells from stresses associated with type 1 diabetes while moderately decreasing insulin secretion in response to high glucose. Further, we show that β cell-specific deletion of the major α subunit isoform, *Scn9a*, mimics and mediates these effects, and can protect mice from type 1 diabetes if induced after weaning. Our findings add support to the concept that a form of excitotoxicity at high glucose concentrations plays a role in type 1 diabetes and suggest that use-dependent Na^+^ channel inhibitors could be viable as future therapeutics.

The roles of ion channels in β cells have been extensively studied in rodents, however there may be important differences in channel expression and properties between rodent and human β cells (70). These differences can have significant effects on β cell electrical activity and insulin secretion, highlighting the need for further investigation of human β cell ion channels and their regulation in health and disease. In the current study, we found that carbamazepine protected human islet cells in a dose-dependent manner from both cytokine- and ER stress-mediated cell death, while having modest inhibitory effects on insulin secretion. These findings confirm that carbamazepine’s previously demonstrated protection from cell death of MIN6 (37) cells and mouse β cells (19) can be translated to human β cells. Despite previous reports that carbamazepine may act via modulation of K_ATP_ channel trafficking (71), structurally related and unrelated Na^+^ channel inhibitors had similar effects and the enhancement of Na^+^ channel activity by veratridine had converse effects on both cell survival and glucose-stimulated insulin secretion. Therefore, the effects of carbamazepine in human β cells are likely primarily mediated by Na^+^ channels. If K_ATP_ channels are involved in these effects of carbamazepine, it must be through a mechanism that also involves SCN9A.

Our work supports the concept that suppression of excitotoxity in the context of hyperglycemia is mechanistic target for type 1 diabetes prevention and therapeutic intervention. Consistent with the ability of Na^+^ channel inhibition to protect neurons (72), previous findings from our lab suggested that reducing voltage-gated Na^+^ channel activity with carbamazepine protects β cells from apoptosis (19). In the current study, oxcarbazepine also reduced cell death but did not reduce glucose-stimulated insulin secretion. We found that other Na^+^ channel inhibitors, spider toxin ProTxII and ambroxol, were only protective in the context of ER stress-mediated cell death and did not significantly affect glucose-stimulated insulin secretion in human islets. These differences may be explained by Na^+^ channel selectivity, as spider toxin ProTxII is more selective for *Scn9a* (73) while ambroxol is more selective for *Scn10a* (74), while carbamazepine has been shown to selectively target *Scn3a* and *Scn9a* (75). The protective effects of carbamazepine can be mimicked by its analogues or other structurally-unrelated drugs, without significant effects on insulin secretion.

Our study adds to the body of knowledge on the physiological and pathophysiological roles of Na^+^ channels in β cells. It was previously believed that Na^+^ channels are more active and relevant in human, canine, and porcine β cells than in rodent cells (14). However, our findings reinforce that Na^+^ channels in mouse β cells should not be disregarded. This is consistent with other studies demonstrating that Na^+^ currents are present in mice (76) and in β cell lines (42). Conditional *Scn9a* deletion in β cells using an *Ins2* promoter-driven Cre transgenic mouse found reduced Na^+^ current density and a non-significant trend towards decreased insulin secretion at 10 mM glucose (40). However, these studies have the caveat that heterozygous controls were used rather than wildtype controls with the normal complement of *Scn9a* channels (40). Here, we show that the *Scn9a* channel plays a key role in insulin secretion at 15 mM glucose, which is probably above the normal physiological range but may be pathophysiological. It has previously been shown that the inhibitory effects of tetrodotoxin on insulin secretion also increase with at higher glucose concentration (77). *Scn9a*’s effect on insulin secretion at high glucose levels are likely coupled to parallel changes in Ca^2+^ signaling. Prolonged exposure to high glucose (i.e. glucotoxicity) and β cell hyperexcitability can cause impaired insulin secretion (78; 79) and has been linked to both type 1 diabetes and type 2 diabetes pathogenesis (80–83). In this context, it is interesting that the *Scn9a* knockout β cells generated using the *Ins1*^Cre^ mouse model and the *Ins1*-Cre AAV8 virus exhibited different electrophysiological behaviour, although there a was an increase in action potential firing at low and moderate glucose in both models.

*Scn9a*^βKO^ cells from NOD mice were protected from ER-stress and cytokine-induced death. Of note, we did not observe a further improvement in cell survival of *Scn9a*^βKO^ islets when carbamazepine was added. These findings demonstrate that the pro-survival effect of carbamazepine is primarily mediated through the *Scn9a* channel. Similarly, the addition of carbamazepine to *Scn9a-*deficient β cells did not further reduce insulin response to glucose. These findings seem to suggest that reduced excitability and the subsequent reduction in insulin secretion protects β cells from stress. Single-cell RNA sequencing of β cell specific *Scn9a*^βiKO^ islets showed gene expression patterns consistent with reduced transcriptional and translational function, along with altered mitochondrial activity and enriched transcriptional regulatory pathways. *Scn9a* loss not only alters electrical activity but is also associated with broad transcriptional reprogramming consistent with a reduced cellular workload, which we have shown is associated with stress resilience and survival in mouse (84; 85) and human (86) β cells.

*Scn9a*^βHET^ mice displayed an exaggerated reduction of glucose-stimulated insulin that was not associated with the same protective effect observed in the homozygous *Scn9a*^βKO^ mice. The exaggerated phenotype was associated with an increase in insulitis, islet apoptosis, as well as a concomitant reduction in insulin area. A 70% reduction in β cell mass is sufficient to cause diabetes in NOD mice (87). It is possible that *Scn9a*^βHET^ phenotype is a product β cell dysfunction caused by inappropriate glucose signaling and impaired distal exocytotic machinery, exacerbated by hyperglycemia. These differences could be explained by a compensatory pathway activated in *Scn9a*^βKO^ mice which remains inactivated in *Scn9a*^βHET^ mice. Bulk RNA sequencing showed differentially expressed genes in *Scn9a*^βKO^ islets that were not observed in *Scn9a*^βHET^ islets. The exaggerated phenotypes associated with *Scn9a*^βHET^ mice observed in our study suggest that from previous studies may need to be re-evaluated. Past work has assigned a relatively minor functional role to *Scn9a* in β cells (40), however those studies used *Scn9a* heterozygous mice as controls, and our results suggest that this control might be inappropriate.

A major finding of our study was that post-weaning, β cell specific *Scn9a* deletion using AAV8*Ins1*Cre, was associated with protection from type 1 diabetes. These results differ from the observations made in *Ins1*^Cre^ mice. These inconsistencies are attributed to the replacement of a single *Ins1* allele and the associated changes in insulin dosage (6; 27; 88). While the AAV8*Ins1*Cre mouse model addresses the limitations associated with insulin dosage, it still relies on Cre to create gene conditional knockouts and as a result is subject to possible protective effects of Cre expression in NOD (27). However, diabetes incidence rates in our WT AAV8*Ins1*Cre control were comparable to wildtype controls with no AAV8*Ins1*Cre and previously reported in WT NOD mice (89). This suggests that the delivery of Cre via AAV8 virus at the 6–7-week time point does not protect against diabetes in these studies. Perhaps the packaging in AAV or the timing when Cre is introduced alters its protective effects. It has previously been reported that the timing of which coxsackievirus is introduced to NOD mice yields the opposing effects on diabetes development (90), by prevention at early exposure and accelerating at late.

The present study establishes *Scn9a* as the likely mediator of the β cell protective and anti-diabetic effects of carbamazepine (19; 20; 37). It will be important to verify clinically that carbamazepine does not have deleterious effects on glucose homeostasis in the context of type 1 diabetes. Carbamazepine administration in people living with epilepsy has not been reported to affect blood glucose levels (91). We found that *Scn9a-*deficient NOD mice did not demonstrate dysfunctional glucose tolerance. Although, these findings suggest that there may be a low risk associated with administration of carbamazepine to patients in a pre-diabetic phase, careful consideration should be made to evaluate the potential impact of reduced insulin secretion under glycemic condition in the context of carbamazepine treatment. In summary, our observations suggest that Na^+^ channels are a promising candidate for future development of therapeutics that aim to protect β cells. A repurposed drug like carbamazepine that can enhance β cell survival in type 1 diabetes may have a shortened path to the clinic.

## Supporting information

All supplemental figures

## Author Contributions

P.O.^†^ co-conceived experiments, conducted experiments, analyzed data and wrote the manuscript. A.G.G.^†^ conducted experiments, analyzed data and wrote the manuscript revisions. X-Q.D. designed, performed, and analyzed electrophysiological studies. N.S.H. and S.P. conducted *in vivo* and *in vitro* experiments. W.G.S. conducted insulin quantification experiments. Y.H.X. performed bioinformatic analysis and data visualization of RNAseq. J.A.Z. performed bioinformatic analysis and data visualization of scRNAseq. E.P. helped conduct scRNAseq experiments. C.M.J.C. helped conduct scRNAseq experiments. H.C. supervised bioinformatic analysis and data visualization. S.S. supervised *in vivo* work. J.K. conducted experiments, supervised work, and edited manuscript. A.G.E. interpreted data and edited the manuscript. P.E.M. supervised work, edited manuscript. J.D.J.* conceived studies, edited manuscript, and is the guarantor of this work and, as such, had full access to all the data in the study and takes responsibility for the integrity of the data and the accuracy of the data analysis.

## Acknowledgements

We thank members of the Johnson and MacDonald labs, as well as members of the JDRF Centre of Excellence for helpful discussions. We also thank Dr. Yike Xie and Dr. Joan Camuñas-Soler (Gothenburg, Sweden) for their valuable guidance with single-cell RNA sequencing analysis. We thank the Human Organ Procurement and Exchange (HOPE) program and Trillium Gift of Life Network (TGLN) for their work in procuring human donor pancreas for research, and James Lyon, Nancy Smith and Dr. Jocelyn Manning Fox (Alberta) for their efforts in human islet isolation. We especially thank the organ donors and their families for their kind gift in support of diabetes research.

