## Supplementary material for "Pharmacological or genetic inhibition of *Scn9a* protects human and mouse beta cells while dampening insulin secretion in type 1 diabetes": All supplemental figures

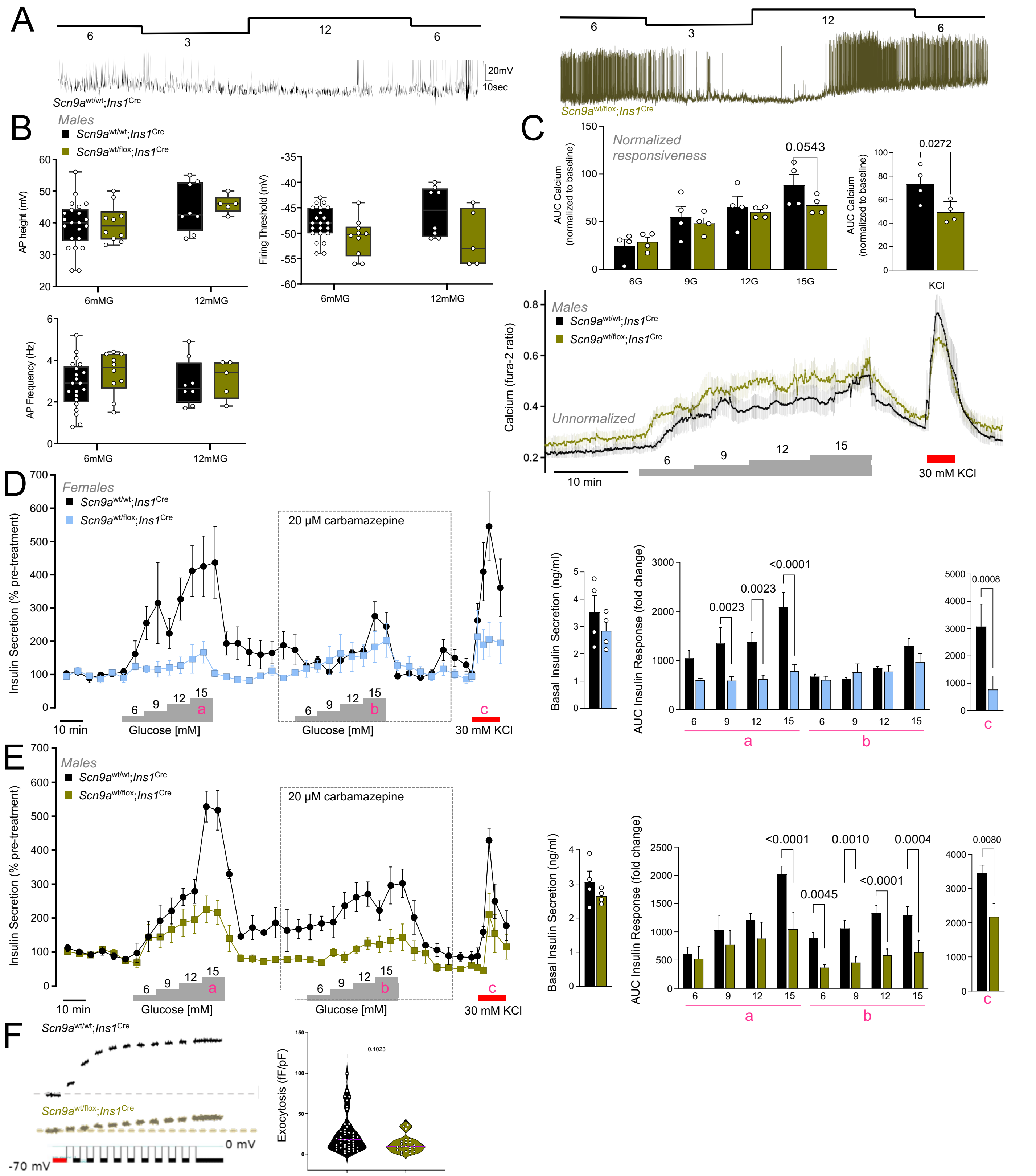


**Figure S1. Heterozygous *Scn9a* knockout mice have reduced glucose stimulated insulin secretion. (A)** Representative membrane potential recording from β cells exposed to 6, 3, and 12 mM glucose. Traces were obtained under the same experimental conditions used for analyses in (B). **(B)** Action potential (AP) frequency, threshold, and height were measured during the initial 6 mM period and subsequent 12 mM glucose period in β cells. Electrophysiology experiments were done in 18-week-old male mice; wildtype (black) and heterozygous (brown). **(C)** Changes in the 340/380 nm Fura-2 ratio measured *Scn9a*-deficient dispersed β cell perifused with Krebs-Ringer bicarbonate buffer containing 3, 6, 9 and 15 mM glucose (grey) or 30 mM KCl (red). Insets AUC of KCl or individual glucose ramp phase normalized to baseline (3 mM glucose) (n = 4-5). **(D,E)** Glucose dependent insulin secretion from 100 whole mouse islets, with or without 20 mM of carbamazepine (dotted line, b) or 30 mM KCl (red line, c) (Normalized to 3 mM glucose baseline ). Insets AUC of KCl or glucose ramp phase. (n = 6). **(F)** Exocytosis measured as increases in cell membrane capacitance by whole-cell patch clamp at 5mM glucose. (n = 3-5). **(C)** AUC data were evaluated using 2-way ANOVA with multiple comparison (mixed models) with Dunnett correction for multiple comparisons. **(F)** Nested one-way ANOVA. Control datasets are identical to those used in corresponding main figures and are shown here for direct comparison.

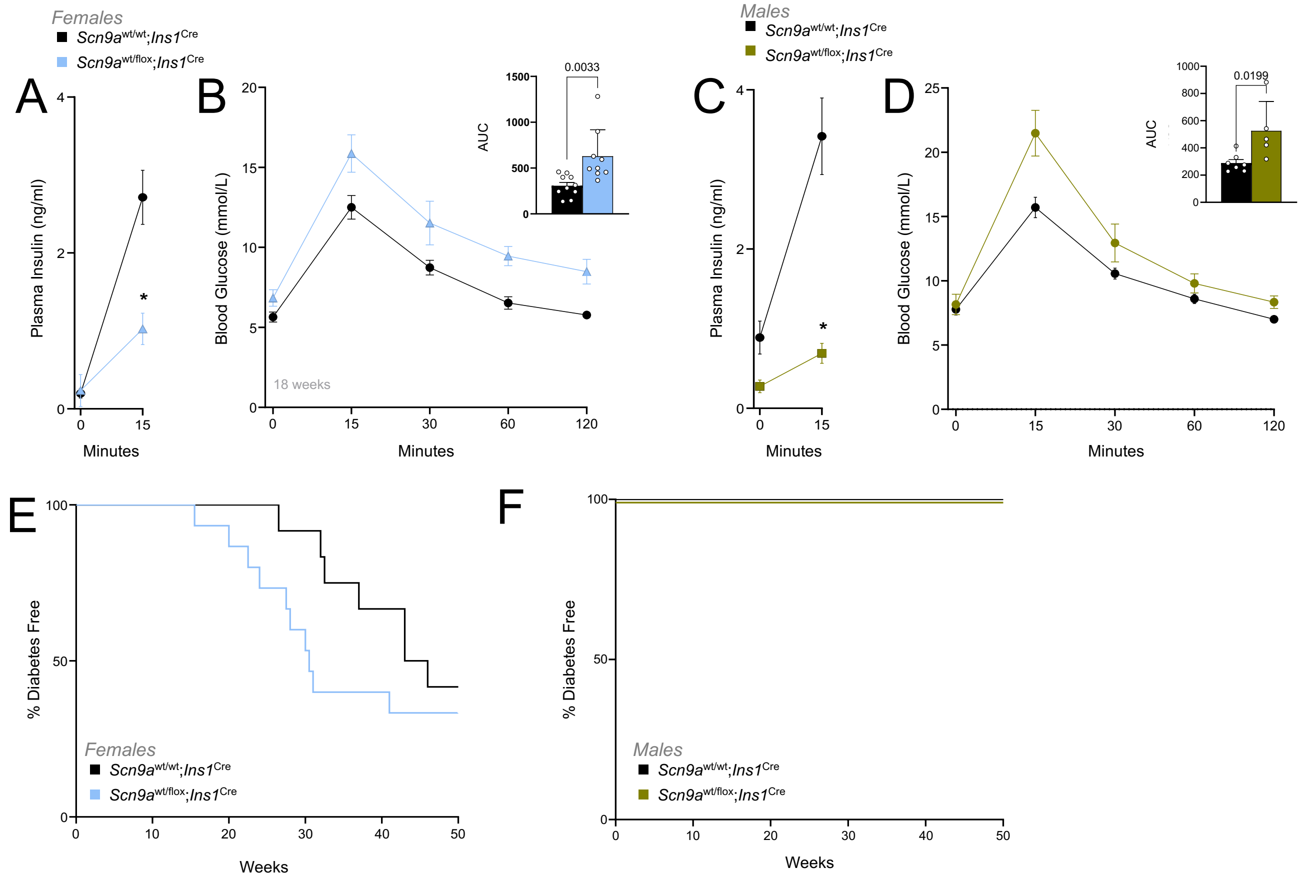


**Figure S2. Impaired glucose tolerance in heterozygous *Scn9a* knockout mice. (A-D)** Insulin secretion and glucose tolerance after intraperitoneal injection (IP) with 2 g/kg of glucose in 18-week-old NOD mice. **(A,C)** Blood was collected at 0 (baseline) and 15 minutes for *in vivo* insulin secretion. **(B,D)** Blood glucose was measured as 0, 15, 30, 60 and 120 minutes after injection. Insets represent area under the curve (AUC) (n = 9). **(E,F)** Kaplan-Meier plot denoting diabetes incidence from male and female NOD mice (n = 12-13). Survival analysis was performed using Log-rank (Mantel-Cox) test. (mean ± SEM, p value shown). Wildtype in black, female in blue, and male mice shown in brown. *p<0.05. **(A-D)** Data were evaluated using 1-way ANOVA. Control datasets are identical to those used in corresponding main figures and are shown here for direct comparison.

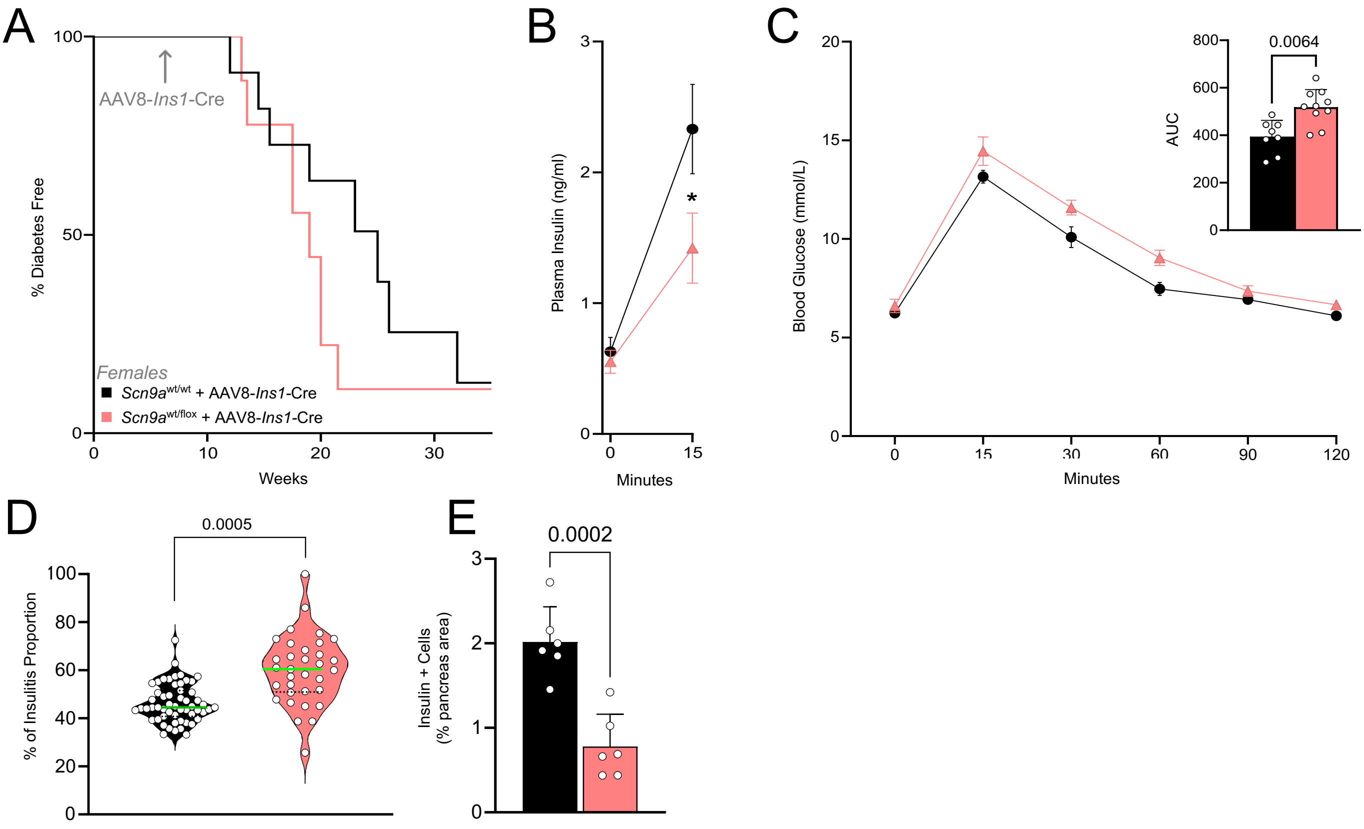


**Figure S3. Heterozygous *Scn9a* knockout mice is not protected from diabetes. (A)** Kaplan-Meier plot denoting diabetes incidence in female NOD wildtype and heterozygous *Scn9a* knockout mice (n = 11-15). Mice were administered 1 × $10^{12}$ VGP AAV8-*Ins1*-Cre via intraperitoneal injection (IP) at 6-7 weeks as indicated by arrow. Survival analysis was performed using Log-rank (Mantel-Cox) test. **(B,C)** *In vivo* insulin secretion and glucose tolerance tests in NOD mice by intraperitoneal injection (IP) with 2 g/kg of glucose at 12-weeks of age. **(B)** Blood was collected at 0 (baseline) and 15 minutes for *in vivo* insulin secretion. **(C)** Blood glucose was measured as 0, 15, 30, 60 and 120 minutes after injection. Inset represents area under the curve (AUC) (n = 9-10). **(D)** Violin plot showing the insulitis proportion using the insulin stain scoring model for 12-week-old NOD mice. **(E)** β-cell area was quantified via insulin staining in 12-week-old mice NOD mice (n = 6). Wildtype mice shown in black and heterozygous knockout mice in red. *p<0.05. **(D)** Data were evaluated using Nested one-way ANOVA. **(A-C and D)** One-way ANOVA. Control datasets are identical to those used in corresponding main figures and are shown here for direct comparison.

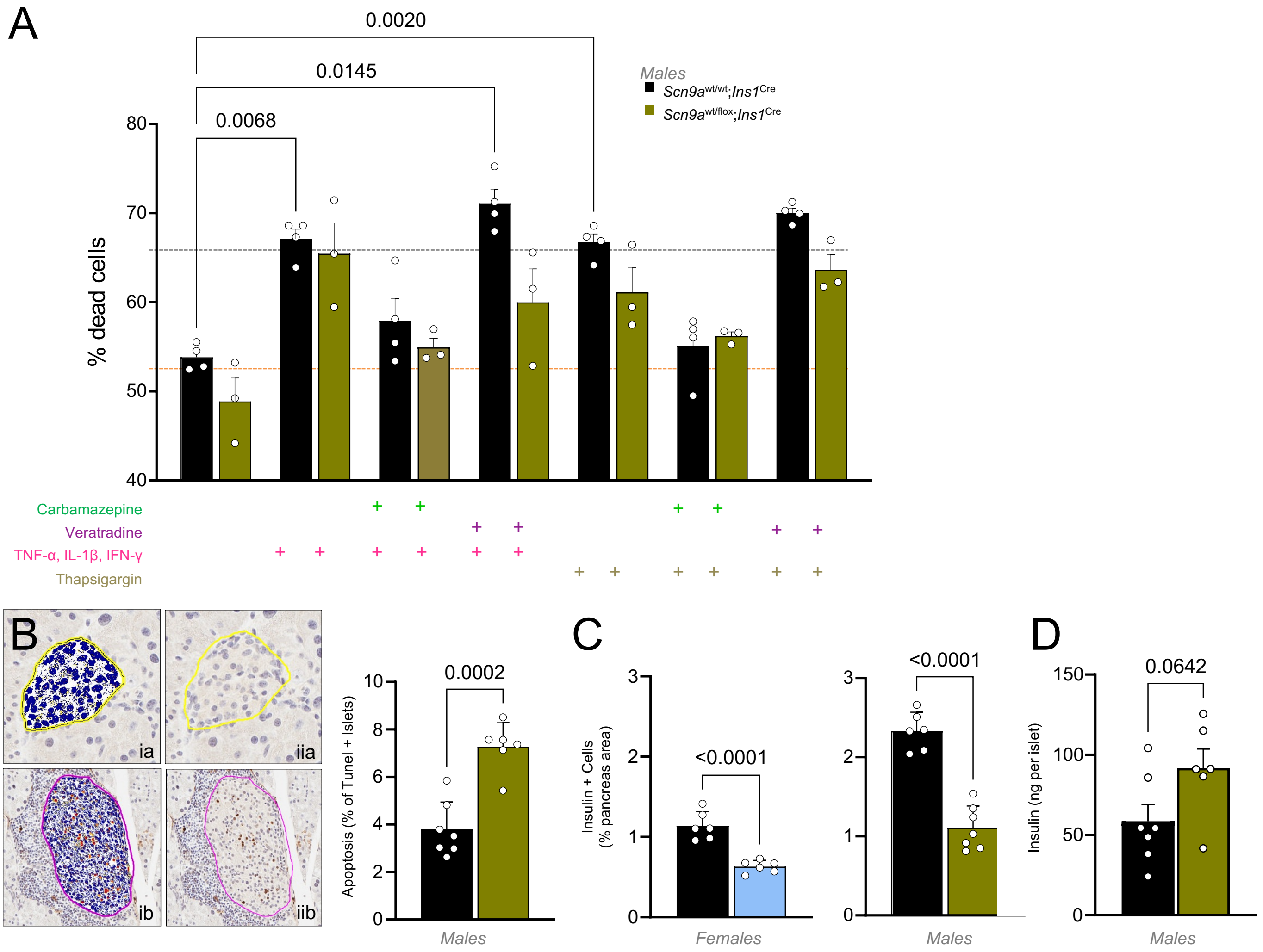


**Figure S4. Heterozygous *Scn9a* knockout mice have increased islet apoptosis and are not protected from ER- and cytokine-induced cell death. (A)** Dispersed islet cells from male, wildtype and heterozygous *Scn9a* knockout mice were seeded onto 384-well plate and stained with Hoechst and propidium iodide (PI), then treated with either a cytokine cocktail or 1 μM thapsigargin, in combination with increasing concentrations of $Na^+$ channel inhibitor carbamazepine, $Na^+$ channel enhancer veratridine or dimethyl sulfoxide (DMSO) control. Cells were imaged with ImageXpress Micro and the percentage of PI-positive (PI+) cells was quantified. Corresponding bar graph shows cell death at 72h time point**. (B)** Representative apoptotic cell death images used for quantification via TUNEL staining (i represent after analysis and ii before, while a-b represent increasing severity of insulitis), in pre-diabetic NOD mice (n = 9). Representative control images shown here are the same as those used in corresponding main text figure. **(C,D)** β-cell area was quantified via insulin staining in NOD mice (n = 6-7). Insulin content in whole islets from male NOD mice (n = 5-7) **(D)**. All experiments were carried out in 18-week-old mice. **(A)** Data were evaluated using 1-way ANOVA with Šídák's multiple comparisons test. **(B-D)** One-way ANOVA. Control datasets shown in panels A-F are the same as those presented in the corresponding main figures. Control datasets are identical to those used in corresponding main figures and are shown here for direct comparison.
